# The receptor kinase SRF3 coordinates iron-level and flagellin dependent defense and growth responses in plants

**DOI:** 10.1101/2021.04.26.441470

**Authors:** Matthieu P. Platre, Santosh B. Satbhai, Lukas Brent, Matias F. Gleason, Magali Grison, Marie Glavier, Ling Zhang, Min Cao, Christophe Gaillochet, Christian Goeschl, Marco Giovannetti, Balaji Enugutti, Marcel von Reth, Ruben Alcázar, Jane E. Parker, Grégory Vert, Emmanuelle Bayer, Wolfgang Busch

## Abstract

Iron is critical for host-pathogen interactions. While pathogens seek to scavenge iron to spread, the host aims at decreasing iron availability to reduce pathogen virulence. Thus, iron sensing and homeostasis are of particular importance to prevent host infection and part of nutritional immunity. While the link between iron homeostasis and immunity pathways is well established in plants, how iron levels are sensed and integrated with immune response pathways remain unknown. We identified a receptor kinase, SRF3 coordinating root growth, iron homeostasis and immunity pathways via regulation of callose synthase activity. These processes are modulated by iron levels and rely on SRF3 extracellular and kinase domain which tune its accumulation and partitioning at the cell surface. Mimicking bacterial elicitation with the flagellin peptide flg22 phenocopies SRF3 regulation upon low iron levels and subsequent SRF3-dependent responses. We propose that SRF3 is part of nutritional immunity responses involved in sensing external iron levels.

## Introduction

Iron is a critical micronutrient for all living organisms. While iron is very abundant in the Earth’s crust, its bioavailability is low. Organisms have evolved efficient iron uptake mechanisms that include a variety of membrane-associated uptake systems that absorb iron unbound or bound to iron-binding molecules. Mammals acquire iron mainly through the glycoprotein transferrin while bacteria, fungi and plants have evolved diverse systems that include siderophores, which are small, high-affinity diffusible secondary metabolites that chelate Fe^3+^ from the surrounding environment (Kramer et al., 2020). In plants, Graminaceae species employ plant specific siderophores while non-Graminaceae such as *Arabidopsis thaliana* depend on an iron reduction-based uptake strategy (Kobayashi and Nishizawa, 2012).

During pathogen attack, iron is at the nexus of host-pathogen interaction as both organisms compete for this metal. Pathogens scavenge iron from the host through siderophore secretion while the host aims to sequester iron to prevent pathogen virulence. Thus, host external iron sensing and internal iron homeostasis regulation are of particular importance to prevent pathogen infection, and are part of the first line of defense called nutritional immunity (Cassat and Skaar, 2013).

In mammals, two receptors, Transferrin Receptor 1 and 2 (TfR) which bind extracellular transferrin-associated iron, play a major role in regulating external iron sensing and homeostasis. Upon host-pathogen interaction, bacterial siderophores outcompete the host iron-bound to transferrin, which in turn leads to a loss of iron triggering independent local and systemic responses in the host (Ganz and Nemeth, 2015). Locally, the loss of iron induces TfR endocytosis and intracellular iron storage via ferritins. Systemically, TfR activation triggers stimulation of the BMPR complex to increase the expression of iron uptake genes (Ganz and Nemeth, 2015). The latter response is intertwined with defense pathway since the inflammatory Interleukine-6 pathway directly interacts with the BMPR complex to regulate iron uptake genes (Ganz and Nemeth, 2015). In *Drosophila melanogaster*, Transferrin-1 was recently shown to activate NF-κB, toll and immune deficiency immunity pathways, thereby mediating nutritional immunity through the control of intracellular iron partitioning (Iatsenko et al., 2020).

Although flowering plants do not contain TfR in their genomes (Bai et al., 2016), iron homeostasis and defense responses are linked (Verbon et al., 2017). Here, FERRETINS (FER) and NATURAL RESISTANCE-ASSOCIATED MACROPHAGE PROTEIN (NRAMPs) were shown to be involved in iron sequestration upon pathogen attack (Deák et al., 1999; Segond et al., 2009). Moreover, the metal transceptor IRON-REGULATED TRANSPORT 1 (IRT1) is critical to mount efficient defense responses (Aznar et al., 2014). Transcriptional signatures of *Pseudomonas simiae* WCS417 and long-term iron deficiency in leaves display an overlap of about 20%, among these genes, the transcription factor MYB DOMAIN PROTEIN 72 (MYB72) plays a role at the interface of both signaling pathways (Dinneny et al., 2008; Zamioudis et al., 2015). Recently, a protein effector from the foliar pathogen *Pseudomonas syringae* was shown to disable a key iron homeostasis regulator, the E3 ligase BRUTUS (BTS), to increase apoplastic iron content and promote colonization (Xing et al., 2021). Finally, the presence of the microbial siderophore, deferrioxamine (DFO), affects the transcriptional landscape of iron homeostasis and immunity genes, suggesting a role for siderophores in mediating nutritional immunity (Aznar et al., 2014). While the link between iron deficiency and immunity is well documented in plants, the mechanism by which iron concentrations are sensed, and how they impinge on iron homeostasis, defense and growth pathways are unknown. Here, we identify the leucine-rich repeats receptor kinase STRUBBELIG RECEPTOR KINASE 3 (SRF3) through *Arabidopsis thaliana* natural root growth variation under low iron levels using a genome wide association study (GWAS). We find that root growth is rapidly reduced upon encountering low iron levels, that *SRF3* is required for this response, and at the same time modulates root iron homeostasis. The regulatory capacity of SRF3 is dependent on its kinase and extracellular domains. Both domains are required for SRF3 partitioning between the plasmodesmata and the so-called bulk PM where it acts as a negative regulator of callose synthases and is degraded upon low iron conditions in both sub-populations. We further establish that SRF3 is a molecular link between responses to low external iron levels and bacterial defense responses, as SRF3 is required to mediate root immune response to the flagellin peptide flg22 by the same mechanisms used for its response to low iron conditions. Our work uncovers a close coordination of responses to low iron levels and immunity pathways and indicates that *SRF3* is located at the nexus of both pathways, thereby constituting a key player in plant nutritional immunity.

## Results

### SRF3 is a regulator of iron homeostasis genes and root growth under low iron levels

Genome wide association studies (GWAS) for root growth rate under low iron levels revealed multiple significantly associated single nucleotide polymorphisms (SNPs) (Figure S1A-B and Spreadheet S1). The most significant association was observed on chromosome 4 in close proximity to the genes AT4G03390 *(STRUBBELIG-RECEPTOR FAMILY 3, SRF3)* and AT4G03400 (*DWARF IN LIGHT 2*, *DFL2*) (Figures 1A).To identify potential causal genes at this locus, we obtained Col-0 T-DNA mutant lines for these genes (Figures S1C-D) and quantified the root growth response after three days exposure to low iron levels. While the *dfl2* T-DNA mutant lines responded similarly to wildtype (WT) to low iron levels, *srf3* T-DNA lines displayed a significantly decreased root growth response compared to WT when exposed to different iron levels using the iron chelator Ferrozine (−FeFZ; 100, 50, 10μM; Figures 1B-C, S1E-G). Moreover, *srf3* mutants showed a slight reduction in their root growth rate in iron sufficient conditions compared to WT and responded similarly to WT to iron excess conditions (Figures 1C and S1G-H). Overall, our data show that *SRF3* is required for an appropriate root growth response to low iron levels.

**Figure 1.**
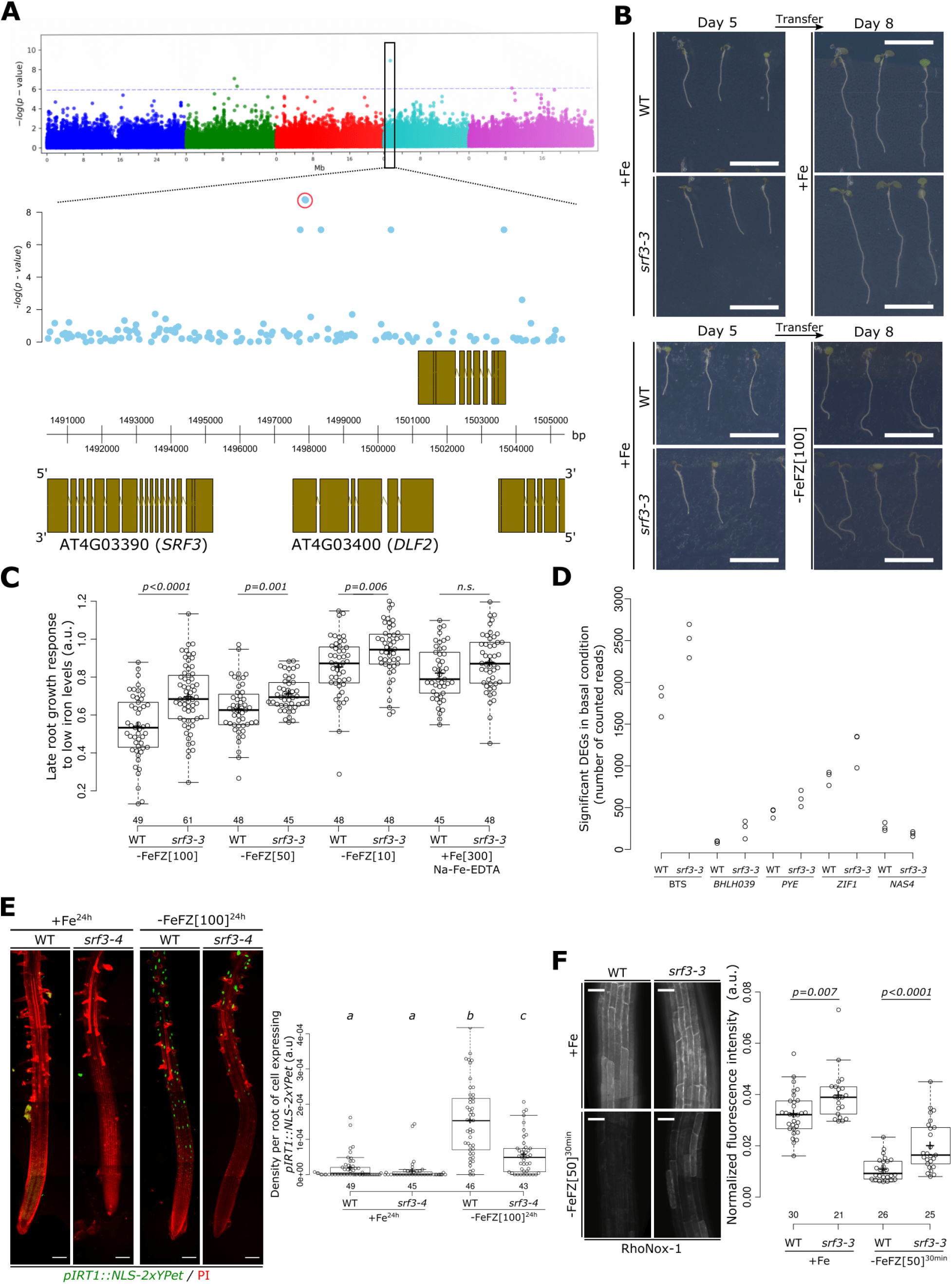
SRF3 regulates root growth and iron homeostasis upon low iron conditions. **(A)** Upper panel: Manhattan plot for GWA mapping of the root growth rate day 4-5 of natural accessions grown under low iron conditions. The horizontal dash dot line corresponds to a 5% false discovery rate threshold. Black box indicates the significantly associated SNP that is in proximity to *SRF3.* Lower panel: Magnified associations in the *SRF3* region with gene models. **(B)** Representative images of 5 days old seedlings of WT and *srf3-3* under iron sufficient medium for 5 days (left panel) and then transferred to iron sufficient media (+Fe; upper right panel), or to low iron medium (−FeFZ 100μM, lower right panel) and grown for 3 days. Scale bars, 1 cm. **(C)** Boxplots of late root growth response to different iron levels (−FeFZ 10,50,100μM or Na-Fe-EDTA 300μM) in WT and *srf3-3* seedlings [two-ways student test (p<0.05), n.s. non-significant]. **(D)** RNAseq read counts of differentially expressed iron homeostasis genes in roots of WT and *srf3-3* in iron sufficient conditions. **(E)** 5 days old seedlings stained with propidium iodide (PI; red channel) expressing *pIRT1::NLS-2xYPet* (green channel) in WT and *srf3-4* on sufficient (+Fe) or low (−Fe) iron medium and the related quantification [one-way ANOVA follows by a post-hoc Tukey HSD test, letters indicate statistical differences (p<0.05)]. Scale bars, 100μm. **(F)** Confocal images of 5 days old seedlings stained with RhoNox-1 in WT and *srf3-3* on sufficient medium (+Fe; upper panel) or low iron medium (ferrozine 50μM, 30min; lower panel) and related quantification [Independent two ways student test (p<0.05)]. Scale bars 50μm.

To explore the function of *SRF3* on iron homeostasis genes, we performed RNAseq on roots from two independent *srf3* alleles and WT under iron sufficient growth conditions. Several key iron homeostasis regulators *(BTS, BHLH039, PYE)* and iron compartmentalization-related genes that are involved in iron transport to the vacuole *(ZIF1)* were upregulated in *srf3* mutants while a key iron distribution transporter involved in iron shoot-to-root partitioning was downregulated *(NAS4;* Figure 1D and Spreadheet S2). Consistent with a mis-regulation of iron responsive genes, the transcriptional reporter line of the low iron inducible iron transporter *IRT1 (pIRT1::NLS-2xYPet)* in *srf3-4* mutant showed a decreased activation after 24 hours under low iron (Figure 1E). In line with a misregulation of iron homeostasis, *srf3* mutants accumulate more iron compared to the WT, thereby phenocopying *bts-1* and *opt3-2,* two iron homeostasis mutants known to accumulate ectopic iron (Figures 1F and S2A-C; Hirayama, 2018; Mendoza-Cózatl et al., 2014; Selote et al., 2015). Importantly, the increased iron levels in *srf3-3* do not stem from increased iron content in the seeds since the iron localization was not altered in *srf3* mutant seeds compared to WT but different from *vit-1* embryos that are known for misdistribution of iron (Kim et al., 2006; Figure S2D). Taken together, these results indicate that *SRF3* is a post-embryonic regulator of iron homeostasis genes.

Next, we investigated the allelic variation at the *SRF3* locus and analyzed accessions according to the pattern of the four top marker polymorphisms associated with the growth response under low iron conditions. The four resulting groups of accessions were haplogroup A that grows slowly on low iron medium and the haplogroups B, C, D that grow faster (Figures S3A-C). While the haplogroup A and haplogroup B differed from several candidate polymorphisms including a larger deletion in the promoter region (Figure S3A), they do not show any significant differences in *SRF3* transcript level accumulation under low iron levels (Figure S3D-E). These results highlight that *SRF3* allelic variation does not lead to obvious changes in *SRF3* transcript levels in bulk root tissue, showing no correlation with the observed variation of root growth rates in low iron conditions. Taken together, our data show that *SRF3* is a negative root growth regulator under low iron levels and is involved in the post-embryonic regulation of iron homeostasis, a function which might be independent of its expression levels.

### The early growth response to low iron is dependent on SRF3 protein levels at the plasma membrane

*SRF3* encodes a gene belonging to the protein family of leucine-rich repeats receptor kinases (LRR-RKs) which are known to be involved in early signal transduction (Hohmann et al., 2017). We hypothesized that *SRF3* might mediate a novel, immediate root response to changes in external iron levels. Using live-light transmission microscopy for 12 hours, we found that low iron levels elicited a significant decrease of root growth after 4 hours and that this response was abolished in *srf3-3* (Figures 2A, S4A and Movie S1-2). The cause of this unresponsiveness in *srf3* is partly explained by a lack of cell elongation decrease upon low iron conditions (Figures S4B-F). To further investigate the role of *SRF3* in regulating root elongation under low iron, we monitored the response of *SRF3* transcription and SRF3 protein abundance using transcriptional and translational reporter lines. A SRF3-2xmCHERRY fusion construct driven by its own promoter fully complemented the *srf3-3* root growth defect under low iron levels, showing the functionality of the construct (Figure S1F and S4G). The transcriptional reporter line revealed that the *SRF3* promoter is active in the differentiation and elongation zones, and to a lesser extent in the transition zone (Figures 2B). Surprisingly, the SRF3 fluorescent protein fusion was detected mainly at the PM in the apical and basal meristem and to a lesser extent in the transition, elongation and differentiation zones (Figures 2B). We confirmed this finding in Landsberg *erecta* (Ler) WT background using a GFP tag fused to the respective full genomic fragment (Figure S4H-L). We reasoned that the SRF3 protein and/or transcript might be cell-to-cell mobile, or that the *SRF3* transcript might expressed transiently in the meristematic cells. Analysis of numerous root tips showed that some roots expressed the *SRF3* transcriptional reporter in the meristematic zone, a finding backed up by single cell sequencing data, however, we could not exclude the alternative hypothesis (Figure S4M-P; Denyer et al., 2019). Overall, *SRF3* is constantly transcribed and translated in the transition-elongation zone and transiently or only in a subset of cells in the meristematic zone.

**Figure 2.**
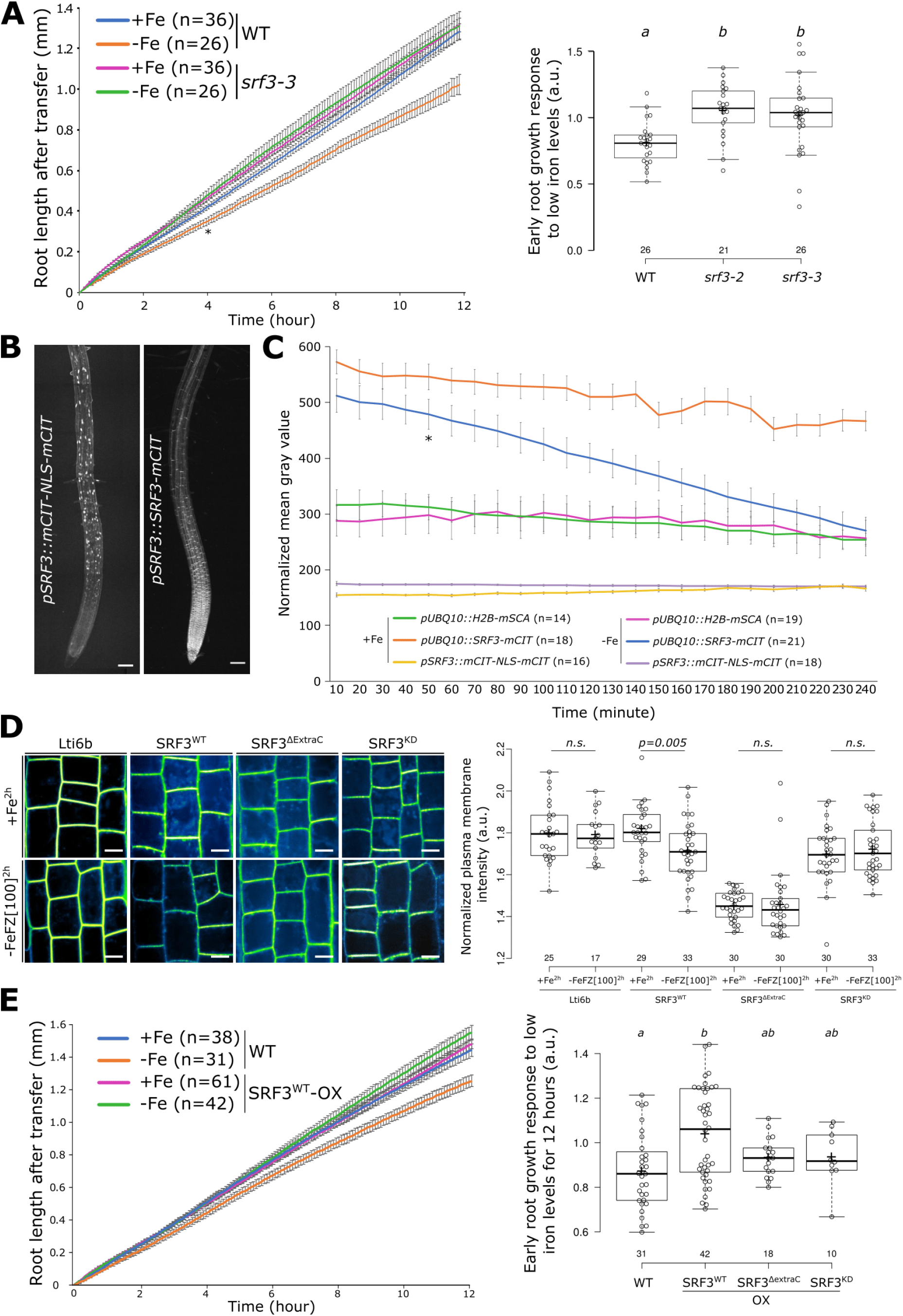
SRF3 regulates early root growth response and undergoes for degradation through its extracellular domain and kinase activity under iron deficiency. **(A)** Graph showing time lapse of the root length of WT and *srf3-3* under sufficient (+Fe) and low (−Fe) iron medias [error bars: SEM; Asterix: significant difference between WT in +Fe and −Fe conditions according to a mixed effect model (*p<0.05*)] and the related quantification including the *srf3-2* mutant [ANOVA with post-hoc Tukey test; Letters: statistical differences (p<0.05)]. **(B)** Confocal images of root tips of 5 days old seedlings expressing *pSRF3::mCITRINE-NLS-mCITRINE* and *pSRF3::SRF3-mCITRINE.* Scale bars, 100μm. **(C)** Graph representing the fluorescence intensity in the root tip of the indicated protein fusions under sufficient (+Fe) and low (−Fe) iron medias [Asterix: significant difference between +Fe and −Fe for *pUBQ10::SRF3-mCITRINE* according to a mixed effect model (*p<0.05*)]. **(D)** Confocal images of 5 days old seedling expressing *p35s::Lti6b-GFP*, *pUBQ10::SRF3^WT^-mCITRINE*, *pUBQ10::SRF3*^ΔExtraC^*-mCITRINE, pUBQ10::SRF3^KD^-mCITRINE* under sufficient (+Fe, 2h) and low iron levels (−FeFZ 100μM, 2h) and the related quantification [two-ways student test (p<0.05), n.s.: non-significant]. Scale bars 10μm. (**E)** Graph showing time lapse of the root length of WT and *SRF3^WT^-OX* under sufficient (+Fe) and low (−Fe) iron medias [error bars: SEM] and related quantification including SRF3^ΔExtraC^ and SRF3^KD^ [ANOVA with post-hoc Tukey test; Letters: statistical differences (p<0.05)].

We next tested whether SRF3 protein abundance or *SRF3* transcription are altered in response to low iron conditions in the transition-elongation zone. While the signal intensity in the *SRF3* transcriptional reporter line did not differ between the two iron regimes, similarly to the control line H2B-mSCARLET, the fluorescent signal intensity in a reporter line in which SRF3 was driven by *UBIQUITIN10* promoter (SRF3^WT^) or its native promotor significantly decreased at the PM under low iron treatment compared to the Lti6b-GFP control line (Figures 2D and S5A-B). Time lapse analysis showed that a signal decrease was recorded after 50 minutes in SRF3^WT^ but not in the other lines (Figures 2C, Movies S3, S4 and S5). We next set out to dissect the role of the functional domains of SRF3 for this process and generated a truncated version of SRF3 in which the extracellular domain had been removed (SRF3^ΔExtraC^) and a kinase dead version, containing a mis-sense mutation in a critical residue in the catalytic ATP binding pocket (SRF3^KD^, Figures S5C-D). While the functional SRF3 protein levels were decreased (SRF3^WT^) after two hours of exposure to low iron conditions, this was not observed for the SRF3^ΔExtraC^ or SRF3^KD^ lines (Figure 2D). This shows that both, the extracellular cellular domain and kinase activity are required to mediate the decrease of SRF3 protein at the PM in response to low iron levels.

We then investigated whether SRF3 levels control early root growth rate under low iron conditions. Surprisingly, much like SRF3 loss of function, constitutive expression of SRF3 abolished the early root growth response to low iron levels (Figures 2E). However, we observed an opposite effect in *srf3* mutant and *SRF3^WT^* overexpressing plants during the late response to low iron (Figure S5E). Although this complex response is yet to be fully explained, we used this property to interrogate SRF3 domain functions. To do so, we investigated the early growth response of the overexpressing lines of *SRF3^ΔExtraC^* (*pUBQ10::SRF3^ΔExtraC^-mCITRINE)*, *SRF3^KD^* (*pUBQ10::SRF3^KD^-mCITRINE)* to low iron conditions. For both early and late low iron growth responses, we observed that roots of *SRF3^ΔExtraC^* and *SRF3^KD^* presented a phenotype close to WT, while the SRF3^WT^ version overexpressing line was hyposensitive or hypersensitive respectively (Figures 2E and S5E). Altogether, our results suggest that the root growth response to low iron conditions requires a fine regulation of SRF3 protein accumulation at the PM, which is dependent on the extracellular and kinase domains. These findings are consistent with a model in which SRF3 senses early apoplastic signals associated with iron depletion through its extracellular domain and transduces the signal(s) intracellularly to modulate root growth via its kinase activity.

### SRF3 resides in two subpopulations at the plasma membrane which are both decreased under low iron conditions

During the analysis of SRF3 expression, we had noticed its enrichment at the PM with a apical-basal localization in punctate foci but also along the entire PM, referred to as bulk PM (Figures S6A and S6C). We tested the role of its extracellular domain and kinase activity for specifying its heterogenous distribution by calculating the standard deviation of the mean intensity (SDMI) along the apical-basal side of PM using SRF3 truncated and point mutant versions. We found that compared to the WT version, SRF3^ΔExtraC^ only associates with the bulk PM since we observed a decrease of the SDMI (Figure 3A) while the removal of the kinase domain (SRF3^ΔKinase^) did not lead to SDMI changes (Figures S5C and S6B). This indicated that the extracellular domain is necessary and sufficient to drive SRF3 into the PM-associated foci. Surprisingly, the standard deviation of SRF3^KD^ fluorescent signal was significantly lower compared to SRF3^WT^ (Figures 3A), suggesting a role of the kinase activity in SRF3 partitioning. We then investigated the role of SRF3 functional domains upon low iron levels. For SRF3^WT^, a decrease of SDMI and polarity upon exposure to low iron conditions were observed, indicating a loss of SRF3-associated punctate structures while such reduction was not observed in the control line, LTI6b-GFP (Figure 3A and S6C). Performing the same experiment with SRF3^ΔExtraC^ and SRF3^KD^ revealed no significant difference upon low iron levels compared to the control condition (Figure 3A). This points to a role of the extracellular domain and a requirement for an active SRF3 kinase in the removal of SRF3 from the foci upon exposure to low iron levels. Taken together, our data show that SRF3 has a dual localization at the plasma membrane, in punctuated structures and the bulk PM that is controlled by the extracellular domain and the kinase activity. Finally, upon exposure to low iron levels, SRF3 seems to become less associated with the punctuated foci which also relies on its functional domains.

**Figure 3.**
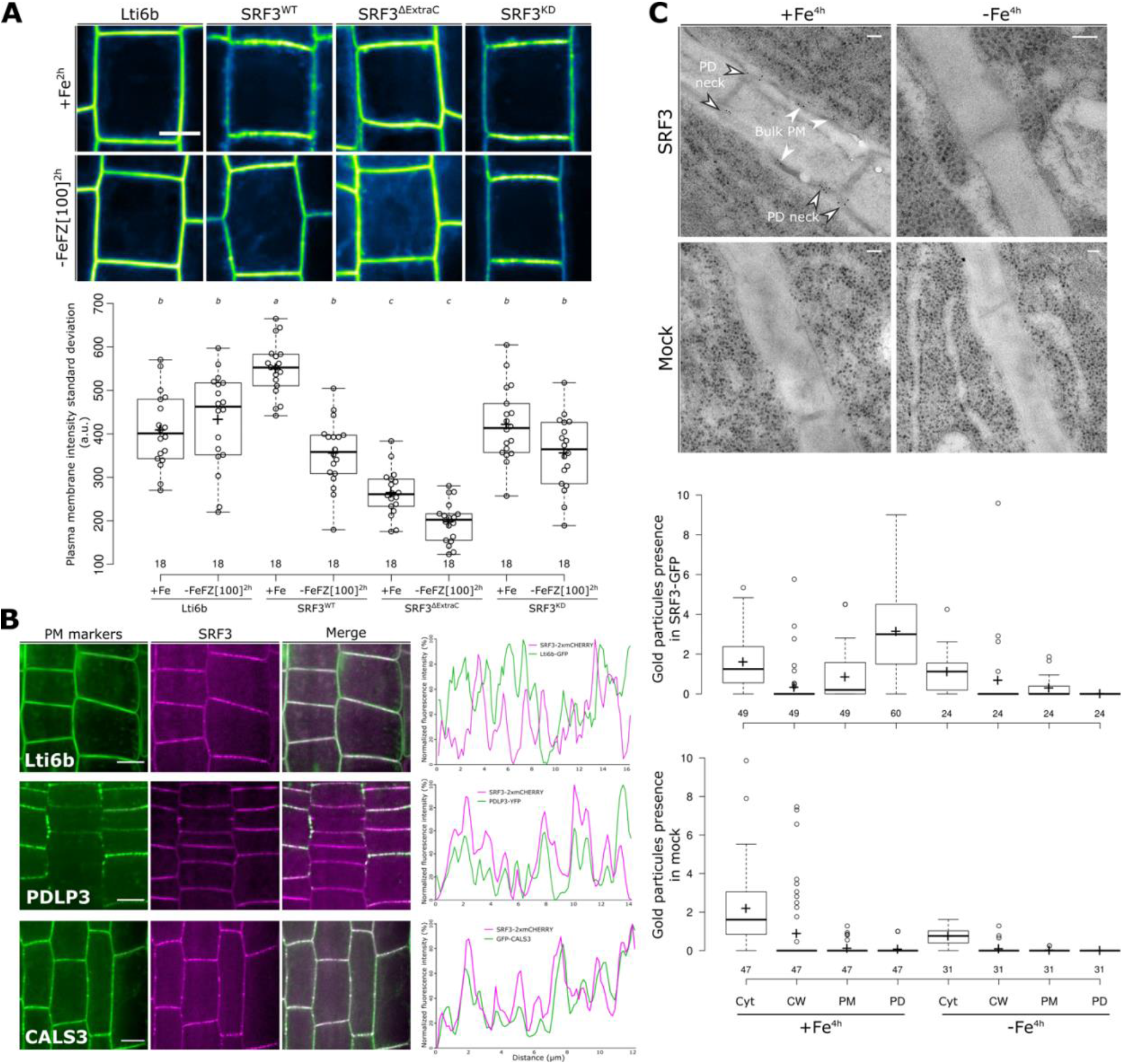
SRF3 co-exists in two subpopulations at the plasma membrane which decrease under low iron levels. **(A)** Confocal images of 5 days old seedling expressing *p35s::Lti6b-GFP*, *pUBQ10::SRF3-mCITRINE, pUBQ10::SRF3^WT^-mCITRINE*, *pUBQ10::SRF3*^ΔExtraC^*-mCITRINE, pUBQ10::SRF3^KD^-mCITRINE* under iron sufficient (+Fe, 2h) or low iron (−FeFZ 100μM, 2h) and the related quantification [ANOVA with post-hoc Tukey test; Letters: statistical differences (p<0.05)]. Note that the pictures have been pseudo-colored to emphasize changes in polarity and localization in the punctuated foci which does not reflect the proper fluorescence intensity. Scale bars, 10μm **(B)** Confocal images of 5 days-old seedlings co-expressing, *p35s::Lti6b-GFP, pPDLP3-PDLP3-YFP, 35s::CALS3-GFP*, left, with *pUBQ10::SRF3-2xmCHERRY*, middle and the relative merge. Red line on the left image indicates where the scan line has been traced. Scale bars, 10μm. Right panel: graphs showing the signal intensity in both channel on the apical basal part of the cell. **(C)** Micrograph of immune gold with plant expressing *pSRF3::SRF3-GFP* (SRF3) and the relative control in *Ler* background under sufficient (+Fe, 4h) and low (−Fe, 4h) iron medias and the related quantification. Cyt, cytosol; CW, cell wall; PM, plasma membrane; PD, plasmodesmata. Scale bars, 100nm

We then addressed the nature of the PM-associated punctuated structures. Analysis of the intensity distribution profile at the PM apical-basal sides of SRF3 fluorescent reporter in the background of PM structure marker lines revealed a specific co-localization of SRF3 with plasmodesmata-associated proteins CALS3 and PDLP3 but not the general PM marker Lti6b (Figure 3B). Moreover, SRF3 co-localized with signals from Aniline blue staining that stains β-1,3- glucan, which are particularly enriched in plasmodesmata (Figure S6D). This strongly indicated that SRF3 localizes or is in close vicinity to the plasmodesma. To characterize and confirm SRF3 subcellular dynamics at higher resolution, we conducted immunogold labeling electron microscopy of the *pSRF3::SRF3-GFP* line using an anti-GFP antibody. In standard conditions, SRF3 signal was localized at the bulk PM and to the plasmodesmata and more specifically to the plasmodesmatal neck region, and was removed not only from plasmodesmata but also the bulk PM under low iron conditions (Figure 3C). As a recent report had shown that some plasmodesmata-associated receptor kinases have a fast and reversible association between bulk PM and plasmodesmata under abiotic stress, which alters their diffusion rates within the PM (Grison et al., 2019), we estimated SRF3 diffusion via fluorescence recovery after photobleaching (FRAP). We found that a decrease of iron levels did not change SRF3 diffusion (Figure S6E), indicating that the decrease of SRF3 is not accompanied by a change in its partitioning. Taken together, our data indicate that SRF3 is associated with the bulk PM but also highly enriched at the neck of the plasmodesmata, in an extracellular domain- and kinase activity-dependent manner. Under low iron, SRF3 becomes depleted from these two subpopulations, a process which is dependent on both SRF3 functional domains.

### Early lack of iron mediates SRF3-dependent callose deposition without affecting cell-to-cell movement

Immunogold-labeling electron microscopy suggested that SRF3 is particularly concentrated at the plasmodesmata neck. This region is highly enriched in sterols, which are required for protein targeting to this specific subregion to regulate plasmodesmata function (Grison et al., 2015). Depleting plants expressing SRF3^WT^ of sterols using sterol inhibitors, Fenpropimorph (Fen) and Lovastin (Lova), showed that SRF3 localization is sterols-dependent since a decrease of SRF3 polarity was observed (Figures S7A), suggesting that SRF3 might have a functional role in this plasmodesmata region. The plasmodesmatal neck is critical for regulating cell-to-cell trafficking, as it is where callose turnover is thought to be regulated to determine plasmodesmata permeability (Sager and Lee, 2018). Iron homeostasis depends on long- and local-distance signaling relying on cell-to-cell movement to activate *IRT1* (Durrett et al., 2007; García et al., 2013; Grillet et al., 2018; Khan et al., 2018; Kumar et al., 2017; Vert et al., 2003). We therefore hypothesized that SRF3 might regulate cell-to-cell communication through callose turnover to properly activate *IRT1*. Analysis of signals from immunostaining with a callose antibody indicated that low iron levels trigger callose deposition in the epidermis and cortex cells of WT root tips (Figure 4A). This shows that iron levels influence callose deposition. In *srf3* mutants, we observed an increase of callose even in the basal condition while callose levels were not responsive to iron depleted media compared to WT (Figure 4A). Our data therefore show that early responses to low iron include an increased callose deposition and that SRF3 negatively regulates this process. To corroborate this finding, we used aniline blue to quantify the signal in the epidermis of the root transition-elongation zone. The positive control, a *CALLOSE SYNTHASE 3* (*CALS3)* overexpressing line, which is known to accumulate ectopic callose showed higher signal intensity compared to WT. In agreement with the antibody based findings, low iron rapidly enhanced callose deposition in WT, however, increased callose was not observed in WT when adding the 2-deoxy-d-glucose (DDG), a well-characterized callose synthase inhibitor (Figures 4B and S7B; Han et al., 2014; Huang et al., 2019; Jaffe and Leopold, 1984; Shikanai et al., 2020; Vatén et al., 2011). In *srf3* mutants, no difference in aniline blue signal intensity was observed under iron sufficient conditions while an increase was observed under low iron compared to WT in the same condition (Figure 4B). Although callose immunostaining and aniline blue slightly differed, both experiments suggest that callose is synthesized by callose synthases shortly after exposure to iron deficiency in an SRF3-dependent manner. Consistent with this conclusion, *srf3-2* and *srf3-3* displayed fused LRs and a higher LR density than WT, both of which are traits associated with higher callose deposition (Benitez-Alfonso et al., 2013; Figures S7C-D).

**Figure 4.**
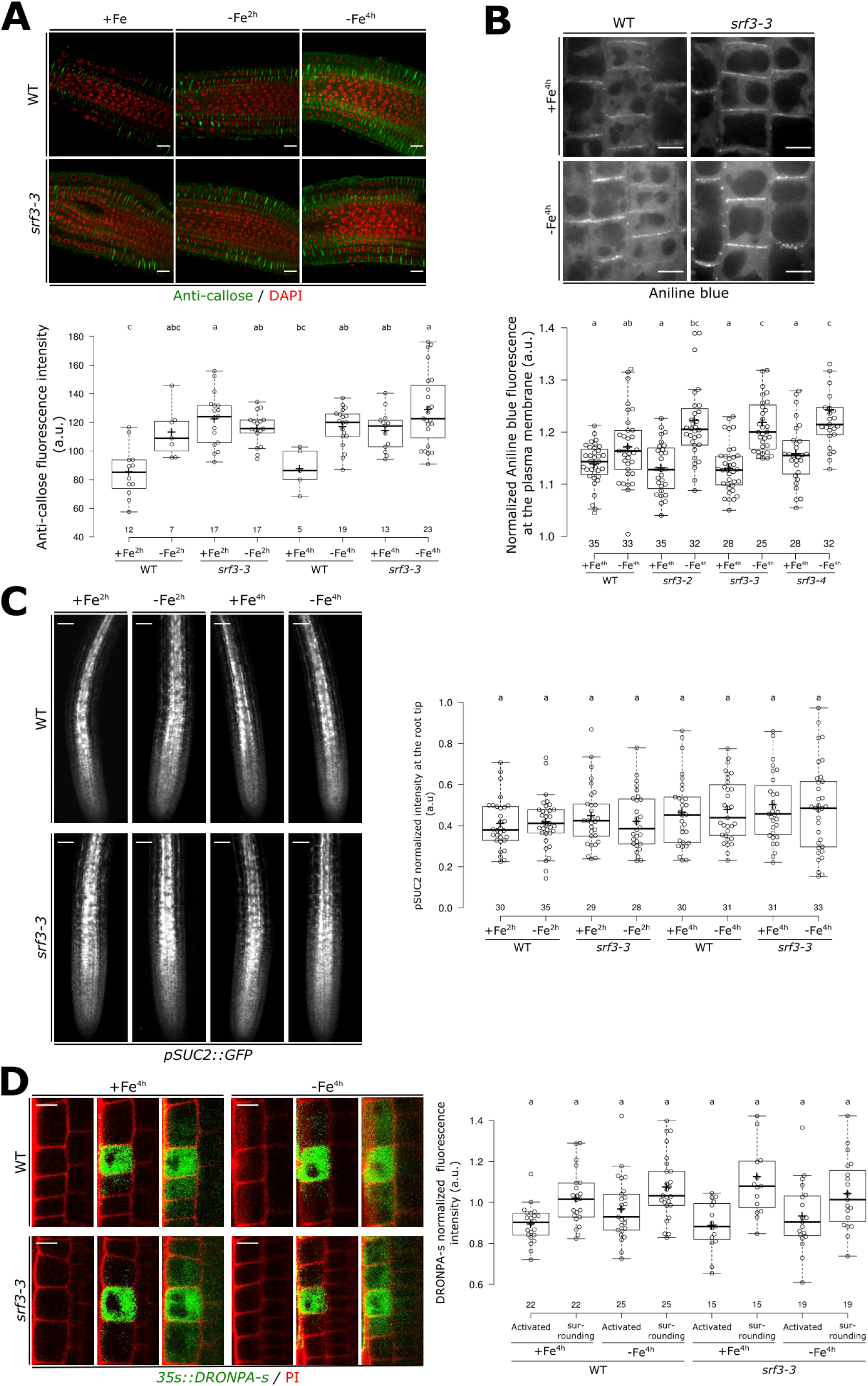
SRF3 is a negative regulator of callose deposition but does not regulate cell-to-cell signaling. **(A)** Confocal images of root of 5 days old seedling stained with callose antibody (green) and DAPI to stain the nucleus (red) under sufficient (+Fe, 2h and 4h) and low (−Fe, 2h and 4h) iron medias and the related quantification [ANOVA with post-hoc Tukey test; Letters: statistical differences (p<0.05)]. Scale bars, 10μm. (**B)** Confocal images of 5 days old seedling stained with aniline blue under sufficient (+Fe, 4h) and low (−Fe, 4h) iron medias in WT and *srf3-3* and the related quantification [ANOVA with post-hoc Tukey test; Letters: statistical differences (p<0.05)]. Scale bars, 10μm. (**C)** Confocal images of 5 days old seedling expressing *pSUC2::GFP* in WT and srf3-3 under sufficient (+Fe) and low (−Fe) iron medias and the related quantification [ANOVA with post-hoc Tukey test; Letters: statistical differences (p<0.05)]. Scale bars, 50μm (**D)** Confocal images of 5 days old seedling expressing *p35s::DRONPA-s* in WT and *srf3-3* under sufficient (+Fe) and low (−Fe) iron medias and the related quantification [ANOVA with post-hoc Tukey test; Letters: statistical differences (p<0.05)]. Scale bars, 10μm.

We next investigated whether callose deposition upon low iron levels modifies cell-to-cell protein movement in a SRF3-dependent manner. We first monitored the ability of GFP expressed in companion cells using *pSUC2::GFP* to diffuse to the surrounding cells through the plasmodesmata, as previously established (Benitez-Alfonso et al., 2013; Nicolas et al., 2017; Vatén et al., 2011a). Surprisingly, no difference in the GFP signal distribution between WT and *srf3-3* root tips from plants grown on iron sufficient and low iron containing media was observed (Figure 4C). To corroborate this observation, we photoactivated DRONPA-s fluorescent protein in a single root epidermis cell and monitored its spread to the upper and lower surrounding cells (Gerlitz et al., 2018). We noticed a decrease of signal in the activated cell and a concomitant increase in the surrounding cells, resulting from cell-to-cell movement. However and consistent with our *pSUC2::GFP* observations, no difference between conditions and/or genotypes was observed (Figure 4D). Altogether, our results suggest that a decrease of iron levels swiftly leads to SRF3- and callose synthase-dependent modulation of callose deposition. However, this does not generally impede cell-to-cell movement.

### Iron homeostasis and root growth are steered by SRF3-dependent callose synthases signaling

While *IRT1* activation is dependent on *SRF3*, this appears not to rely on a restriction of cell-to-cell movement via callose synthases-mediated callose deposition during the early responses to low iron conditions. We therefore reasoned that *IRT1* regulation might rely on early signaling events that are dependent on callose synthases, or that *IRT1* regulation only occurs at a later stage of the response. We first tested whether *IRT1* is regulated at the time during which SRF3-dependent callose deposition occurs. A 16-hour time lapse analysis of *IRT1* promoter activity indicated that the *IRT1* promoter becomes active during the first hours of low iron conditions while no or little activity was observed in iron sufficient media (Figures 5A, Movie S6 and S7). In *srf3-4* mutant roots, we observed a lower expression of the *IRT1* reporter line upon low iron conditions compared to WT, indicating that early *IRT1* transcriptional activation depends on SRF3 (Figure 5B). Next, we tested whether callose synthases activity was important to activate *IRT1* transcription by inhibiting callose synthases with DDG. The addition of DDG in low iron conditions strongly reduced *IRT1* promotor activation in WT, which was not observed in the *srf3-4* mutant compared to mock conditions (Figure 5B). All together, these observations indicate that *SRF3* likely acts upstream of callose synthases-mediated signaling to ultimately tune the expression of the major root iron transporter IRT1.

**Figure 5.**
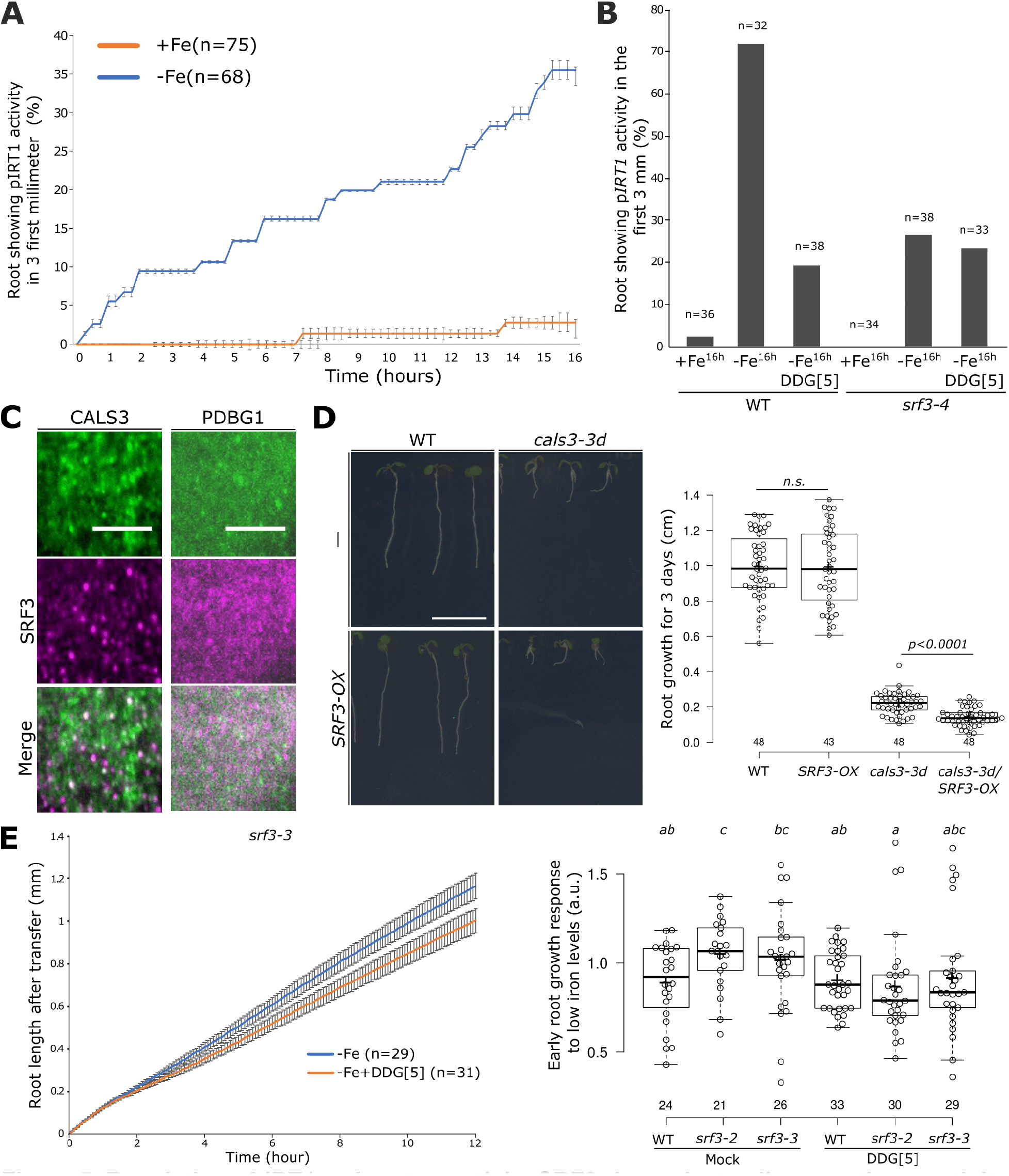
Regulation of *IRT1* and root growth by SRF3-dependent callose synthase activity under low iron levels. **(A)** Graph representing the quantification of *pIRT1::NLS-2xYPet* time lapse analysis under mock (+Fe) and low iron levels (−Fe) [error bars indicate SEM]. **(B)** Graph representing the percentage of root showing *IRT1* promotor activation under sufficient (+Fe, 16h) and low (−Fe, 16h) iron medias in WT and *srf3-4* in presence or absence of DDG. **(C)** Micrographs of 5 days old seedling expressing *35s::GFP-CALS3* (upper) and *UBQ10::SRF3-2xmCHERRY* (middle) and merge channel (lower) in TIRF. Scale bars, 5μm. (**D)** Picture of 9 days old seedling of WT, *cals3-3d*, *pUBQ10::SRF3-mCITRINE* (*SRF3-OX)* and *cals3-3d*x*SRF3-OX* and the related quantification [two-ways student test (p<0.05), n.s. non-significant]. Scale bar, 1cm. **(E)** Graph showing time lapse of the root length of *srf3-3* under low iron (−Fe) medias in presence or absence of DDG and the related quantification including the *srf3-2* mutant [ANOVA with post-hoc Tukey test; Letters: statistical differences (p<0.05); Error bars: SEM].

We next investigated the regulatory interaction of SRF3 and callose synthases by colocalization analysis in roots using dual-color total internal reflection fluorescence (TIRF) revealing that SRF3 was organized in microdomains that partially colocalized with CALS3 but not with the β-1,3-glucanases reporter PdBG1 known to negatively regulate callose deposition (Figure 5C). To test these interactions genetically, we crossed *SRF3-OX,* which does not present any root growth defects, with a mutated version of CALS3 (*cals3-3d*) whose activity is up to 50% higher and subsequently accumulates more callose, resulting in shorter roots than in WT (Vatén et al., 2011a). The double homozygous lines of *SRF3-OX*x*cal3-3d* showed a further decrease of root growth compared to the *cals3-3d* single mutant (Figures 5D and S7E), indicating a genetic interaction of SRF3 and CALS3. To test whether the observed phenotype was due to a specific genetic interaction or a more general interaction of increased callose levels and SRF3 overexpression, we used the *35s::GFP-PDLP5* (*PDLP5-OX*) line, which presents similar callose and root growth phenotypes as observed in *cals3-3d* (Lee et al., 2011a; Sager et al., 2020). In this SRF3-OXx*PDLP5-OX* line, the root growth phenotype was indistinguishable from the *PDLP5-OX* line, therefore highlighting the specific genetic interaction of *SRF3* and *CALS3* (Figures 5D and S7E). Next, we set out to further test whether *SRF3* is acting upstream or downstream of CALSs. We reasoned that if CALSs were upstream of SRF3, the inhibition of callose synthase activity would impact SRF3 PM levels. Co-treatment of low iron with DDG did not modify PM-associated SRF3 levels and therefore suggested that callose synthase is downstream of SRF3 (Figure S7F). This finding was corroborated by monitoring the early and late root growth rate of WT and *srf3-3* during the application of DDG and low iron levels as a partial complementation of *srf3-3* root growth phenotype was observed in that condition (Figures 5E and S7G). Overall, our data suggest that SRF3 acts upstream of the callose synthase early-on upon low iron levels to regulate iron homeostasis and root growth.

### SRF3 coordinates iron homeostasis and bacteria elicited immune responses

*SRF3* was originally identified as a genetic locus underlying immune-related hybrid incompatibility in *Arabidopsis* and shown to be involved in bacterial defense-related pathways in leaves (Alcazar et al., 2010). Gene ontology (GO) analysis of root RNAseq data in standard condition and the analysis of the root specific *pCYP71A12::GUS* immune reporter upon treatment with the bacterial elicitor flg22 showed that SRF3 has a similar role in roots (Figure 6A and S8A; Millet et al., 2010). We then investigated the specificity of SRF3’s role by assessing the late root growth responses to different pathogen-associated molecular patterns (PAMPs) and plant-derived damage-associated molecular patterns (DAMPs). *srf3* roots were only impaired in their response to flg22 but not to chitin or AtPep1 compared to WT (Figures S8B-C). Similar to the low iron level response, the flg22 response was already apparent early-on in WT and absent in *srf3* mutants (Figures 6B, S8D-E and Movies S8, S9). To test whether the increased iron content of *srf3* might be related to this response, we analyzed the early and late root growth responses to flg22 of *bts-1* and *opt3-2.* They both responded like WT, indicating that higher iron root content does not generally affect the root growth regulation upon immune response elicitation (Figures S8F-H; Mendoza-Cózatl et al., 2014; Selote et al., 2015). We therefore concluded that the role of *SRF3* in controlling early root growth upon bacterial elicitation is specific and related to its signaling activity.

**Figure 6.**
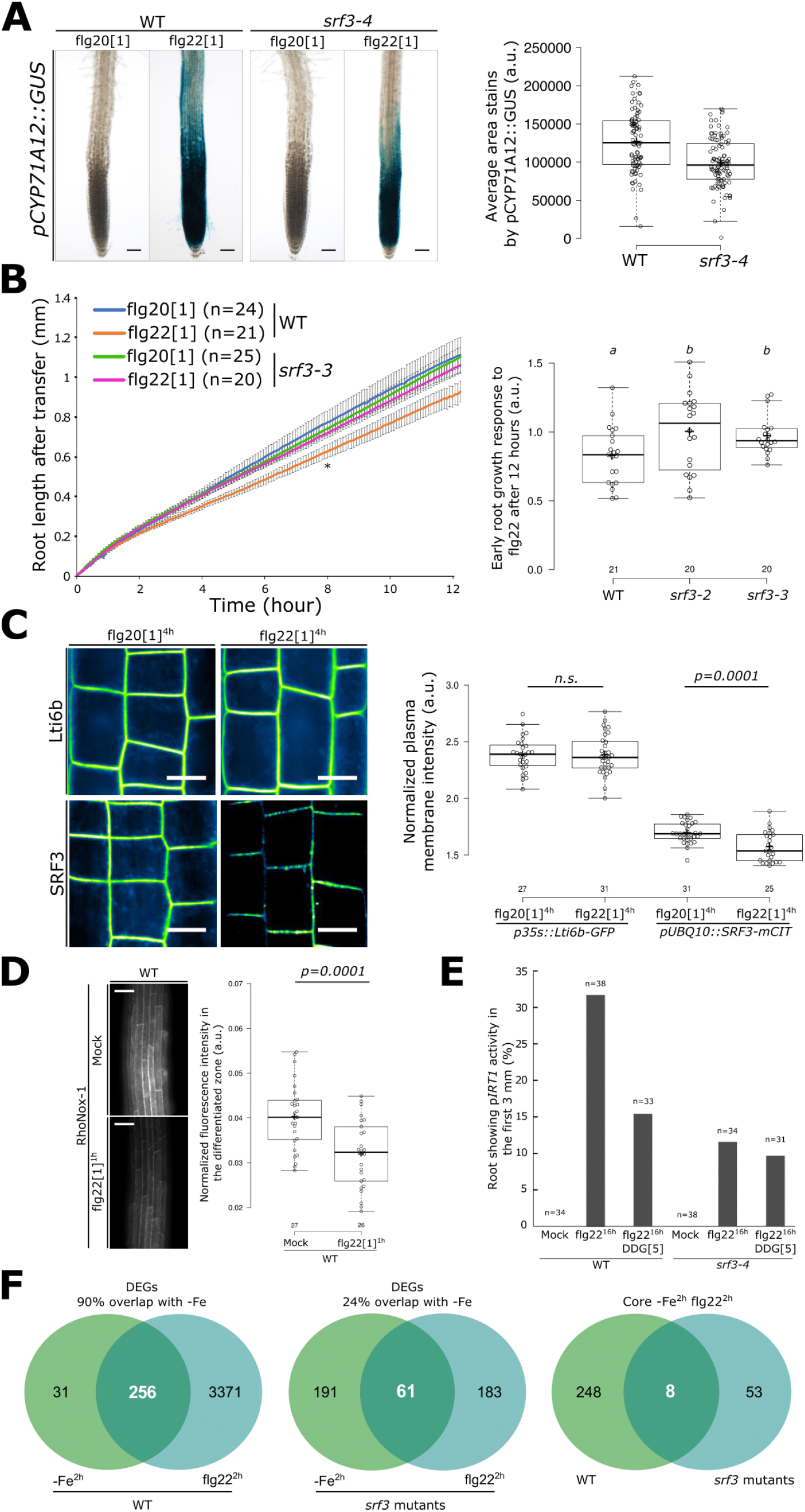
Coordination of bacterial immunity and iron homeostasis signaling pathways by SRF3. **(A)** Pictures of plants expressing *pCYP71A12::GUS* in WT and *srf3-4* under flg20 and flg22 treatment (1μM, 24h) and the related quantification. Scale bars, 50μm. (**B)** Graph showing time lapse of the root length of WT and *srf3-3* under flg20 and flg22 (1μM) [Error bars: SEM; Asterix: significant difference between +Fe and −Fe for the WT according to a mixed effect model (*p<0.05*)] and the related quantification including the *srf3-2* mutant [ANOVA with post-hoc Tukey test; Letters: statistical differences (p<0.05)]. (**C)** Confocal images of 5 days old seedling expressing *p35s::Lti6b-GFP* and *pUBQ10::SRF3-mCITRINE* in flg20 and flg22 (1μM, 4h) and the related quantification [two-ways student test (p<0.05); n.s.: non-significant]. Scale bars, 10μm. **D)** Confocal images of 5 days old seedling stained with RhoNox-1 in WT in mock or flg22 (1μM, 1h) and the related quantification [two-ways student test (p<0.05)]. Scale bars, 50μm. (**E)** Graph representing the percentage of root showing *IRT1* promotor activation under mock and flg22 (1μM, 16h) treatment in WT and *srf3-4* in presence or absence of DDG. (**F)** Venn diagram of differentially expressed genes under low iron levels (−Fe, 2h) and flg22 (1μM, 2h) in WT (left) in *srf3* (middle) and DEGs in both condition between WT and *srf3*.

Because of the similar growth response to low iron levels and flg22 treatment, we hypothesized that the SRF3-dependent root growth regulation to these two stresses might rely on the similar molecular mechanism. Consistent with this idea, we found that upon flg22 treatment, the SRF3 protein displays similar cellular dynamics as observed under low iron conditions (Figure 6C and S8I) while no significant changes of SRF3 transcriptional regulation were observed (Figure S8J). Therefore, SRF3 appears to be a point of convergence between iron and flg22-dependent signaling mediating root growth regulation.

We reasoned that one model explaining this convergence is that flg22 might trigger a transient decrease of cellular iron levels, thus promoting SRF3 degradation. We therefore performed RhoNox-1 staining after 1 hour of flg22 treatment and observed a decrease of fluorescence compared to mock treatment (Figure 6D). This indicates a swift decrease of local iron concentration in roots upon flg22 stimulus. Consistent with a rapid decrease of cellular iron, flg22 treatment rapidly enhanced *IRT1* expression (Figure 6E, S9A and Movie S10). Moreover, mining of publicly available root RNAseq data revealed a broad impact of short-term flg22 treatment on the expression of iron homeostasis genes (Spreadheet S3; Stringlis et al., 2018). We then wondered whether the flg22-triggered iron deficiency responses rely on SRF3-dependent callose synthase activity. Co-treatment of WT roots with DDG and flg22 led to a decrease of *IRT1* promoter activity compared to flg22 treatment alone, while in *srf3-4* this activation was decreased under flg22 compared to WT and insensitive upon co-treatment (Figure 6E). Overall, these data indicate that flg22-dependent *IRT1* activation relies on SRF3-mediated callose synthase signaling as observed for low iron conditions.

Finally, to investigate the extent of *SRF3*-dependent coordination of bacterial immune responses and iron homeostasis, we performed an RNAseq analysis after two hours of exposure to low iron levels or flg22 in *srf3* mutant and WT roots. Strikingly, 90% of the differentially expressed genes (DEGs) in these two conditions overlapped and were up or down regulated in the same manner in WT. Importantly, these DEGs were not associated with a general stress response since none of these common iron and flg22 DEGs were overlapping with those in cold, NaCl and mannitol datasets (Figure 6F, S9B and Spreadheet S4 and S5; Kreps et al., 2002). To further confirm that low iron levels trigger immunity genes, we conducted qPCR for two early markers of flg22-triggered immunity, *FRK1* and *MYB51* that showed a transient activation of the two genes within four hours (Figure S9C; He et al., 2006). To determine how much of this common transcriptional program is coordinated by SRF3, we analyzed the srf3 transcriptome datasets. DEGs in flg22 and iron deficiency in *srf3* mutant roots only overlapped by 24% demonstrating that *SRF3* coordinates a large part of the common transcriptional program that is triggered in response to early response to low iron and to flg22 (Figure 6F). Overall, our work establishes SRF3 as a major coordinator of bacterial immune response and iron deficiency signaling pathways which relies on callose synthase signaling.

## Discussion

Based on a GWAS approach, we have identified an LRR-RK, *SRF3* as a regulator of early root growth responses to low iron conditions. We show that SRF3 transduces signals that lead to a coordinated response of root growth regulation, iron homeostasis and bacterial immunity through its modulation of callose synthase-dependent signaling. Because this is highly reminiscent of nutritional immunity conferred by the TfR mammalian and *Drosophila* systems that sense iron levels and control iron and immune responses, we propose that *SRF3* is instrumental in mediating plant nutritional immunity (Cassat and Skaar, 2013; Iatsenko et al., 2020).

### The root responses to low iron are triggered rapidly and mediated by SRF3 signal transduction

We discovered that root growth is modulated within the first four hours upon exposure to low iron levels looking at earlier time points than usually considered (Figure 2A; Durrett et al., 2007; Hindt et al., 2017; Mendoza-Cózatl et al., 2014; Satbhai et al., 2017)). This early response is SRF3-dependent, exposing this LRR-RK as being a key part of the genetically encoded ability of roots to perceive and transduce low environmental iron levels. A comprehensive SRF3 domain characterization showed that the LRR and the kinase activity are critical not only for its organization at the PM but also to mediate SRF3 decrease-dependent root growth arrest under low iron (Figure 2). In light of other LRR-RK signalling transduction mechanisms, such as those for BRI1 and FLS2 (Belkhadir and Jaillais, 2015; Hohmann et al., 2017; Jaillais and Vert, 2016; Tang et al., 2017), our results lead towards the following model for SRF3 1) the LRR extracellular domains senses a signal that is informative of the early lack of iron, 2) which in turn activates the kinase activity 3) which then triggers decrease of its level at the PM, 4) to regulate early root growth. We also found that the role of SRF3 in transducing low iron levels at an early stage is not restricted to the root growth regulation according to the RNAseq analysis (Figure 1E, 5B and 6). However, we did not provide direct evidence of the involvement of SRF3 kinase activity and LRR in transducing signals to regulate iron homeostasis and bacterial immune pathways. However, this is very likely since SRF3 is known to be part of the phosphorelay upon PAMP immune response (Benschop et al., 2007). Much of the *SRF3* function is tied to the SRF3 signal transduction because no obvious changes in SRF3 transcriptional level in flg22 or low iron treatments were observed (Figures S5A and S8J), and no correlation between the expression level of *SRF3* in accessions that displayed contrasting root growth responses to low iron levels were observed (Figure S3E). Yet, an early or cell-type specific *SRF3* transcriptional regulation cannot be excluded. Altogether, our data indicate that roots perceive external variation of iron rapidly through SRF3-dependent signal transduction to coordinate root signaling pathways.

### Early responses to low iron are mediated by SRF3-dependent callose synthases regulation

We have found that SRF3 acts upstream of iron-induced callose synthases activity to mediate proper signaling. Aniline blue and immunostaining showed that the early low iron response goes along with callose synthase-dependent callose deposition. Even though, these two techniques indicated conflicting results for callose deposition levels in the basal condition in WT and *srf3-3*, which might be explained by technical reasons, both approaches pinpointed SRF3 acts as negative regulator of this process (Figures 4A and 4B). The role of SRF3 acts as an upstream negative regulator of callose synthases is further strengthened by several lines of evidence: SRF3 and CALS3 colocalize in both SRF3 PM subpopulations (Figure 3B and 5C), genetically act in the same pathway and the root growth response to low iron levels of *srf3* mutants is partially complemented upon inhibition of callose synthases (Figure 5B-E). Surprisingly, cell-to-cell movement of proteins were not affected early-on upon low iron levels, despite callose synthases activation and increased callose deposition in the plasmodesmata (Figure 4C-D). However, it is possible that callose deposition might impact later responses since callose deposition-mediated plasmodesmata closure can take hours to days to occur (Cheval et al., 2020; Lee et al., 2011a; Lim et al., 2016; Rutschow et al., 2011; Stonebloom et al., 2012). Another possibility is that callose deposition might have a different function early-on, for instance early ROS signaling which is thought to mediate callose deposition, actually increases cell-to-cell communication in leaves (Fichman et al., 2021). Thus, even though root growth, iron homeostasis and defense signaling can be controlled by cell-to-cell movement of signaling molecules movement, SRF3 dependent regulation of these pathways doesn’t rely on impeding cell-to-cell movement thereby putting the spotlight onto a signaling function of callose synthases. In line with this idea, the double mutant *SRF3-OX*/*cals3-3d* displayed shorter roots compared to *cals3-3d* which should in fact show longer roots if SRF3 was strictly restricted to its repressive role on callose deposition (Figures 5D). Taken together, we have found that SRF3-dependent callose synthase activity is required to regulate early root growth, iron homeostasis and defense signaling pathways under low iron levels, which might dependent directly on callose synthase-mediated signaling rather than impeding cell-to-cell movement.

### *SRF3* mediates early root responses to low iron levels and a bacterial PAMP

We have found that *SRF3*-mediated signaling is at the nexus of the early root responses to low iron and bacterial-derived signal. In fact, RNAseq analysis revealed that early responses to low iron and flg22 are highly similar and largely coordinated by *SRF3* (Figure 6F). The axis of SRF3 and callose synthases is of particular importance for the regulation iron homeostasis genes in both conditions as revealed by monitoring *IRT1* promotor activity (Figure 5B and 6E). The local and swift decrease of iron in the root upon flg22 treatment might be the mechanism that underlies the flg22 dependent activation of iron homeostasis genes (Figure 6D-C). Conversely, the early lack of iron is able to activate the PTI signaling pathways, which is also mediated by SRF3 (Figures 6F and S9C). This activation of PTI signaling upon low iron levels is likely due to SRF3’s role to modulate iron homeostasis which is important to coordinate immune responses (Figures 1D-E and 5B Spreadheet S2; Palmer et al., 2013; Verbon et al., 2017). However, there is an alternative model that cannot be excluded. In this model SRF3 regulates flg22-mediated PTI signaling pathways, which in turn modulates iron homeostasis. This model is in line with the specific SRF3-dependent root growth regulation under flg22, the RNAseq data from *srf3* mutant roots in which PTI-dependent genes are misregulated (Spreadheet S2, S4 and Figure S9D) and experimental data provided in Smakowska-Luzan et al., 2018, based on the extracellular network. Further supporting this hypothesis, we found that the early and late root growth responses of *fls2* mutants, which are impaired in PTI-triggering immunity are decreased under low iron levels (Figure S9E-H).

Altogether, our observations lead to a model in which SRF3 perceives an early lack of iron to modulate iron homeostasis and PTI signaling pathways, however it remains to be investigated which pathway is upstream of the other.

### The interaction of low iron levels and pathogens

During host-pathogen interactions, an early host line of defense is to withhold iron to limit pathogen virulence, which is part of the nutritional immune responses as previously reported in vertebrates and invertebrates (Ganz and Nemeth, 2015; Iatsenko et al., 2020). Eliciting bacterial immune responses triggers a SRF3-dependent decrease of cellular iron levels, showing a conserved principle of this nutritional immune response being present in plants (Figures 1F and 6D). In line with this idea, we have found that the lack of *SRF3* impedes mechanisms relating to the ability of root tissues to withhold iron. For instance, *ZIF1* that is involved in iron storage in the vacuole is upregulated in *srf3* mutants, while *NAS4* that is involved in root-to-shoot iron transport is downregulated (Figure 1D; Spreadheet S2; Haydon et al., 2012; Klatte et al., 2009). In parallel, *NAS4* modulates ferritin accumulation, which is another way for the plant to withhold iron (Koen et al., 2013). Moreover, similar to the nutritional immunity systems described in mammals and *Drosophila melanogaster* that are based on TfR, SRF3 senses the immediate lack of iron which is also relayed to a common signaling pathway linking iron deficiency and immunity responses (e.g. BMPR; Figure 6F; Cassat and Skaar, 2013). Altogether, we therefore propose that SRF3 is a central player in a mechanism that embodies a fundamental principle of nutritional immunity by coordinating bacterial immunity and iron signaling pathways via sensing iron levels.

## Acknowledgements

We thank Y. Belkhadir, B. Lacombe, I. Helariutta and all the Busch lab members for critical discussions. Y. Jaillais for sharing cloning materials, F. Berger for providing H2B in PDONR P1P2, J.B.D Long for providing SV40, Y. Benitez-Alfonso for providing 35s:PdBG1-mCITRINE line, I. Helariutta for providing feedback on the manuscript and sharing 35s:GFP-CALS3 and cals3-3d as well as R. Stadler for providing 35s:DRONPA-s line. J.Y. Lee for providing 35s::PDLP5-GFP line. E. Bayer for providing pPDLP3::PDLP3-YFP and pSUC2::GFP lines. pCYP72A::GUS line was kindly provided by Y. Belkhadir. We thank, T. Zhang and the Salk Biophotonics core team for microscopy advance and assistance in quantification. We thank as well Salk peptide synthesis core especially Jill Meisenhelder and finaly Br. Moussu for thoughtful discussion. This study was funded by the National Institute of General Medical Sciences of the National Institutes of Health (grant number R01GM127759 to W. Busch), a grant from the Austrian Science Fund (FWF I2377-B25 to W. Busch), funds from the Austrian Academy of Sciences through the Gregor Mendel Institute (W.Busch), and start-up funds from the Salk Institute for Biological Studies (W. Busch). M.P. Platre was supported by a long-term postdoctoral fellowship (LT000340/2019 L) by the Human Frontier Science Program Organization. R.A. and J.E.P. were supported by The Max-Planck Society and Germany’s Excellence Strategy CEPLAS (EXC-2048/1, Project 390686111). M.v.R was funded by an IMPRS PhD fellowship. The European Research Council (ERC) under the European Union’s Horizon 2020 research and innovation program (grant agreement No 772103-BRIDGING) to E. Bayer with the EMBO Young Investigator Program to E. Bayer.

## Author contributions

M.P. Platre was responsible of all experiments described in the manuscript except for : qRT-PCR from extreme accessions performed by M. Giovannetti, B. Enugutti did dry seed embryo dissection, Marcel von Reth, R. Alcazar and Jane E. Parker generated and characterized the *pSRF3-SRF3-GFP* line, G. Vert provided pIRT1-NLS-2xYPET line, S.B. Satbhai was involved in phenotyping, GWAS data processing and analysis, and performing *pCYP72A11::GUS* experiment. C. Goeschl performed GWAS data plotting and GUS signal quantification, L. Brent performed the selection and generation of transgenic lines, M.F. Gleason imaged SRF3 reporter lines, M. Cao conducted qRT-PCR for immune genes under iron deficiency, C. Gaillochet and L. Zhang performed RNAseq data analysis, M. Glavier performed SRF3 immuno-gold electron microscopy and M. Grison performed callose immuno-localization. M.P. Platre, S.B. Satbhai and W. Busch conceived the study and designed experiments. M.P. Platre, W.Busch and E. Bayer wrote the manuscript, and all the authors discussed the results and commented on the manuscript.

## STAR METHODS

### KEY RESOURCES TABLE

| REAGENT or RESOURCE | SOURCE | IDENTIFIER |
| --- | --- | --- |
| Antibodies |  |  |
| Anti-callose | Bio-supply | CAT #400-2 |
| anti-GFP | Thermo Fisher Scientific | CAT #A11122 |
| anti-rabbit IgG | Tebu-Bio | CAT #NA.41 |
| Chemicals, peptides, and recombinant proteins |  |  |
| DDG | Tocris | CAT #4515 |
| Flg20 | Salk peptide synthesis core | N/A |
| Flg22 | Salk peptide synthesis core | N/A |
| Ferrozine | Acros Organics | CAT #AC410570010 |
| Na-Fe-EDTA | PhytoTechnology Lab | CAT #E349 |
| Aniline blue | Sigma | CAT #28631-66-5 |
| RhoNox-1 | Goryo Chemical | CAT #GC901 |
| Experimental models: organisms/strains |  |  |
| *A. thaliana*:<br>*Col-0* | NASC | NASC# N1092 |
| *A. thaliana*:<br>*srf3-2* | Alcazar et al., 2010 | NASC #557621 |
| *A. thaliana*:<br>*srf3-3* | NASC | NASC #501389 |
| *A. thaliana*:<br>*srf3-4* | NASC | NASC #878396 |
| A. thaliana:<br>*cals3-3d* | Vatén et al., 2011 | N/A |
| A. thaliana:<br>*pCYP71A12::GUS* | Millet et al., 2010 | N/A |
| A. thaliana:<br>*pIRT1::NLS-2xYPet* | This study | N/A |
| A. thaliana:<br>*p35s::DRONPA-s* | Gerlitz et al., 2018 | N/A |
| A. thaliana:<br>*pUBQ10::H2B-mSCARLET* | This study | N/A |
| A. thaliana:<br>*pSRF3::mCITRINE-NLS-mCITRINE* | This study | N/A |
| A. thaliana<br>*pSRF3::SRF3-GFP/Ler* | This study | N/A |
| A. thaliana:<br>*pSRF3::SRF3-2xmCHERRY/srf3-3* | This study | N/A |
| A. thaliana:<br>*pSRF3::SRF3-mCITRINE/srf3-3* | This study | N/A |
| A. thaliana:<br>*pUBQ10::SRF3-mCITRINE* | This study | N/A |
| A. thaliana:<br>*pUBQ10::SRF3<sup>KD</sup>-mCITRINE* | This study | N/A |
| A. thaliana:<br>*pUBQ10::SRF3<sup>ΔKinase</sup>-mCITRINE* | This study | N/A |
| A. thaliana:<br>*pUBQ10::SRF3<sup>ΔExtraC</sup>-mCITRINE* | This study | N/A |
| A. thaliana:<br>*35s::mCITRINE-PdBG1* | Benitez-Alfonso et al., 2013 | N/A |
| A. thaliana:<br>*35s::GFP-CALS3* | Vatén et al., 2011 | N/A |
| A. thaliana:<br>*35s::PDLP5-GFP* | Lee et al., 2011 | N/A |
| A. thaliana:<br>*pPDLP3::PDLP3-YFP* | Thomas et al., 2008 | N/A |
| Oligonucleotides |  |  |
| For primers used in genotyping, RT-PCR and cloning see Spreadsheet S6, related to STAR Methods | N/A | N/A |
| Recombinant DNA |  |  |
| Empty gateway entry vector: pDONR221 | thermofisher | CAT# 12536017 |
| Empty gateway entry vector: pDONR P2RP3 | thermofisher | CAT# 12537023 |
| Empty gateway destination vector: pB7m34GW | (Karimi et al., 2007) | N/A |
| Empty gateway destination vector: pH7m34GW | (Karimi et al., 2007) | N/A |
| Gateway entry vector (promoter): UBQ10prom/<br>pDONR P4P1R | (Jaillais et al., 2011) | NASC# N2106315 |
| Gateway entry vector (promoter): pSRF3/ pDONR<br>P4P1R | This study | N/A |
| Gateway entry vector (protein): SRF3/ pDONR221 | This study | N/A |
| Gateway entry vector (protein): SRF3<sup>ΔKinase</sup>/<br>pDONR221 | This study | N/A |
| Gateway entry vector (protein): SRF3ΔExtraC/<br>pDONR221 | This study | N/A |
| Gateway entry vector (protein): SRF3KD/<br>pDONR221 | This study | N/A |
| Gateway entry vector (fluorescent protein):<br>H2BnoSTOP/pDONR221 | This study | N/A |
| Gateway entry vector (fluorescent protein):<br>2xmCHERRYnoSTOP/ pDONR P2RP3 | (Platre et al., 2018) | N/A |
| Gateway entry vector (fluorescent protein):<br>mCITRINEnoSTOP/pDONP2RP3 | (Jaillais et al., 2011) | NASC# N2106287 |
| Gateway entry vector (fluorescent protein):<br>mSCARLET/pDONP2RP3 | This study | N/A |
| Gateway destination vector (for plant<br>transformation): pUBQ10::H2B-mSCARLET/<br>pH7m34GW | This study | N/A |
| Gateway destination vector (for plant<br>transformation): pSRF3::SRF3-2xmCHERRY<br>pH7m34GW | This study | N/A |
| Gateway destination vector (for plant<br>transformation): pSRF3::SRF3-mCITRINE/pB7m34GW | This study | N/A |
| Gateway destination vector (for plant<br>transformation): pUBQ10::SRF3-mCITRINE/pB7m34GW | This study | N/A |
| Gateway destination vector (for plant<br>transformation): pUBQ10::SRF3KD-mCITRINE/pB7m34GW | This study | N/A |
| Gateway destination vector (for plant<br>transformation): pUBQ10::SRF3ΔKinase-<br>mCITRINE/pB7m34GW | This study | N/A |
| Gateway destination vector (for plant<br>transformation): pUBQ10::SRF3ΔExtraC-mCITRINE/pB7m34GW | This study | N/A |
| Gateway destination vector (for plant<br>transformation): pUBQ10::SRF3-2xmCHERRY/pH7m34GW | This study | N/A |
| Software and algorithms |  |  |
| BRAT | (Slovak et al., 2014) | N/A |
| Fiji | (Schindelin et al., 2012) | https://fiji.sc/ |
| Matlab_RootWalker | Busch's lab | N/A |
| Macro_PM_intensity | Busch's lab | N/A |
| Macro_RhoNox-1 | Busch's lab | N/A |
| Macro_Match_Align | Busch's lab | N/A |
| Macro_Nuclear_signal_Intensity | Busch's lab | N/A |
| Macro_Nuclear_density | Busch's lab | N/A |

#### Plant materials and growth conditions

For surface sterilization, *Arabidopsis thaliana* seeds of 231 accessions from the Regmap panel (Spreadheet S1) that had been produced under uniform growth conditions were placed for 1 h in opened 1.5-mL Eppendorf tubes in a sealed box containing chlorine gas generated from 130 mL of 10% sodium hypochlorite and 3.5 mL of 37% hydrochloric acid. For stratification, seeds were imbibed in water and stratified in the dark at 4 °C for 3 days. Seeds were then put on the surface of 1X MS agar plates, pH 5.7, containing 1% (w/v) sucrose and 0.8% (w/v) agar (Duchefa Biochemie) using 12-cm x 12-cm square plates. The iron-sufficient medium contained 100 μM Na-Fe-EDTA and the iron-deficient (1XMS iron free) medium contained 300 μM Ferrozine, a strong iron chelator [3-(2-pyridyl)-5,6-diphenyl-1,2,4-triazinesulfonate, Sigma Aldrich](Dinneny et al., 2008). This condition was only used for GWAS. For further experimentation, we used the Fe -sufficient or -free media described in Gruber et al., 2013, with no or a decrease level of Ferrozine, 100, 50 and 10 μM. Using the Gruber et al., 2013 iron-free medium, we add 300 μM of Na-Fe-EDTA to test *srf3* phenotype under iron excess(Gruber et al., 2013). The *srf3-2*, *srf3-3, bts-1, opt3-2*, *fls2-c, fls2-9, vit-1 and cals3-3d* mutant lines are in Col-0 background and were described and characterized(Alcázar et al., 2010; Groen et al., 2013; Higashi et al., 2008; Kim et al., 2006; Mendoza-Cózatl et al., 2014; Selote et al., 2015; Vatén et al., 2011b). The reporter lines, *35s::PdBG1-mCITRINE, 35s::GFP-CALS3, pPDLP3::PDLP3-YFP, 35s::GFP-PDLP5, 35s::Lti6b-GFP, 35s::DRONPA-s, pSUC2::GFP* and *pCYP71A12::GUS* are in Col-0 background and were described and characterized(Benitez-Alfonso et al., 2013; Cutler et al., 2000; Lee et al., 2011b; Millet et al., 2010; Thomas et al., 2008b; Vatén et al., 2011b). The T-DNA insertion lines for SRF3, SAIL1176_B01 (*srf3-4*) and SALK_202843, as well as for *at4g03400,* SAIL_811_C06 (*at4g03400*) were purchased from Nottingham Arabidopsis Stock Center (NASC, Nottingham, United Kingdom). The primers used for genotyping the T-DNA lines are listed below (List of primers, Spreadheet S6). Plants were grown in long day conditions (16/8h) in walk in growth chambers at 21°C, 50uM light intensity, 60% humidity. During nighttime, temperature was decreased to 15°C.

#### Plant transformation and selection

Each construct (see below: “Construction of plant transformation vectors (destination vectors) and plant transformation”), was transformed into C58 GV3101 *Agrobacterium tumefaciens* strain and selected on YEB media (5g/L beef extract; 1g/L yeast extract; 5g/L peptone; 5g/L sucrose; 15g/L bactoagar; pH 7.2) supplemented with antibiotics (Spectinomycin, Gentamycin). After two days of growth at 28C, bacteria were collected using a single-use cell scraper, re-suspended in about 200 mL of transformation buffer (10mM MgCl2; 5% sucrose; 0.25% silwet) and plants were transformed by the floral dipping method(Clough and Bent, 1998). Plants from the Columbia–0 (Col0) accession were used for transformation. Primary transformants (T1) were selected *in vitro* on the appropriate antibiotic/herbicide (glufosinate for mCITRINE, hygromycin for mCHERRY and mSCARLET tagged proteins). Approximately 20 independent T1s were selected for each line. In the T2 generation at least 3 independent transgenic lines were selected using the following criteria when possible: i) good expression level in the root for detection by confocal microscopy, ii) uniform expression pattern, iii) single insertion line (1 sensitive to 3 resistant segregation ratio) and, iv) line with no obvious abnormal developmental phenotypes. Lines were rescreened in T3 using similar criteria as in T2 with the exception that we selected homozygous lines (100% resistant). At this step, we selected one transgenic line for each marker that was used for further analyses and crosses.

#### GWA mapping

231 natural accessions (12 plants/accession were planted) were grown on 1 × MS agar plates containing 300 μM Ferrozine under long day conditions (16 hours light) at 21°C. Plant images were acquired by EPSON flatbed scanners (Perfection V600 Photo, Seiko Epson CO., Nagano, Japan) every 24 hours for 5 days (2 DAG – 6 DAG). Root image analyses and trait quantification were performed using the BRAT software(Slovak et al., 2014). Median root growth rate (n ≥3) values between 4 to 5 days were used for GWA study. For more accuracy, the roots not detected or not germinated were not included in the analyses. GWA mapping was conducted using a mixed model algorithm which has been used previously to correct population structure confounding and SNP data from the 250K SNP chip(Atwell et al., 2010; Brachi et al., 2010; Horton et al., 2012; Kang et al., 2008; Seren et al., 2012). SNPs with minor allele counts equal or greater to 10 were taken into account. The significance of SNP associations was determined at 5% FDR threshold calculated by the Benjamini-Hochberg-Yekutieli method to correct for multiple testing(Benjamini and Yekutieli, 2001). The GWAS peak in proximity of *SRF3* (Figure 1a) contained 4 significant SNPs. By analyzing the unique combinations of these 4 SNPs in the 231 accessions, four groups of haplotypes were defined as Group A, Group B, Group C and Group D.

#### Phenotyping of early root growth responses

Seeds were sowed in +Fe media described in Gruber et al., 2013 and stratified for 2-3 days at 4°C. Five days after planting, about 15 seedlings were transferred to a culture chamber (Lab-Tek, Chamberes #1.0 Borosilicate Coverglass System, catalog number: 155361) filled with – Fe or +Fe medium described in Gruber et al., 2013 or +Fe medium containing flg20 or flg22. Note that the transfer took about 45-60 seconds. Images were acquired every 5 minutes for 12 hours representing 145 images per root in brightfield conditions using a Keyence microscope model BZ-X810 with a BZ NIKON Objective Lens (2X) CFI Plan-Apo Lambda.

#### Phenotyping of late root growth responses

Seeds were sowed in +Fe media described in Gruber et al., 2013 and stratified for 2-3 days at 4°C. Five days after planting, 6 plants per genotype were transferred to four 12×12cm plates in a pattern in which the positions of the genotypes were alternating in a block design (top left, top right, bottom left and bottom right). After transfer, the plates were scanned every 24 hours for 3 days using the BRAT software(Slovak et al., 2014).

#### Quantitative real time PCR

For *SRF3* expression analysis seedlings were grown initially on iron sufficient media (1xMS, 1% w/v Caisson Agar) for 5 days and then shifted to either iron sufficient or low iron (100 uM FerroZine) 1xMS liquid medium. Nylon mesh (Nitex Cat 03-100/44; Sefar) was placed on top of the solidified media to facilitate transfer. Root tissues were collected for RNA extraction 3 hours post transfer by excision with fine scissors. Each biological replicate was constituted by RNA extraction from 30-40 whole roots. Samples were immediately frozen in liquid nitrogen, ground, and total RNA was extracted using the RNeasy Plant Mini kit (QIAGEN GmbH, Hilden, Germany). qRT-PCR reactions were prepared using 2x SensiMix SYBR & Fluorescein Kit (PEQLAB LLC, Wilmington, DE, USA) and PCR was conducted with a Roche Lightcycler 96 (Roche) instrument. Relative quantifications were performed for all genes with the β-tubulin gene (AT5G62690) used as an internal reference. The primers used for qRT-PCR are shown in list of primers (Spreadheet S6).

#### RNAseq

Total RNA was extracted from roots of plants 5 days after germination using RNA protein purification kit (Macherey-Nagel). Next generation sequencing (NGS) libraries were generated using the TruSeq Stranded mRNA library prep kits (Illumina, San Diego, CA, USA). Libraries were sequenced on a HiSeq2500 (Illumina, San Diego, CA, USA) instrument as single read 50bases. NGS analysis was performed using Tophat2 for mapping reads onto the Arabidopsis genome (TAIR10)(Kim et al., 2013), HT-seq for counting reads and EdgeR for quantifying differential expression(Robinson et al., 2009). We set a threshold for differentially expressed genes (Fold change (FC) >2 or FC<-2, FDR<0.01). Genotype x Environment interaction analysis was performed using linear model and type II Anova analyses in R (codes are available upon request). Gene ontology analysis was performed using the AgriGOv2 online tool(Tian et al., 2017). Venn diagrams were generated with the VIB online tool (http://bioinformatics.psb.ugent.be/webtools/Venn/). The plot in figure 3a was generated using the online Revigo software (http://revigo.irb.hr/).

#### Microscopy setup

All imaging experiments except when indicated below, were performed with the following spinning disk confocal microscope set up: inverted Zeiss microscope equipped with a spinning disk module (CSU-X1, Yokogawa, https://www.yokogawa.com) and the prime 95B Scientific CMOS camera (https://www.photometrics.com) using a 63x Plan-Apochromat objective (numerical aperture 1.4, oil immersion) or low resolution 10x lens for time lapse imaging. GFP, mCITRINE, Aniline blue and RhoNox-1 staining were excited with a 488 nm laser (150mW) and fluorescence emission was filtered by a 525/50 nm BrightLine® single-band bandpass filter. mSCARLET, mCHERRY and propidium iodide dyes were excited with a 561 nm laser (80 mW) and fluorescence emission was filtered by a 609/54 nm BrightLine® single-band bandpass filter (Semrock, http://www.semrock.com/). 405 nm laser was used to excite aniline blue and emission was recorded at 480–520 nm with 40x objectives. For propidium iodide, 488nm for excitation and around 600nm was used with 40x objectives. For quantitative imaging, pictures of root cells were taken with detector settings optimized for low background and no pixel saturation. Care was taken to use similar confocal settings when comparing fluorescence intensity or for quantification.

#### FRAP experiment

Fluorescence in a rectangle ROI (50 μm2, 15 μm long), in the plasma membrane region, was bleached in the root optical section by four successive scans at full laser power (150 W) using the FRAP module available on the Zeiss LSM 980 Airyscan 2. Fluorescence recovery was subsequently analysed in the bleached ROIs and in controlled ROIs (rectangle with the same dimension in unbleached area). FRAP was recorded continuously during 90 s with a delay of 0.3 s between frames. Fluorescence intensity data were normalized as previously described (Platre et al, 2019). For visualization, kymographs were obtained using kymograph function in Fiji.

#### TIRF microscopy

Total Internal Reflection Fluorescence (TIRF) Microscopy was done using the inverted ONI Nanoimager from Oxford microscope with 100x Plan-Apochromat objective (numerical aperture 1.50, oil immersion). The optimum critical angle was determined as giving the best signal-to-noise ratio. Images were acquired with about 15% excitation (1W laser power) and taking images every 100ms for 500-time steps.

#### DRONPA-s bleaching and activation

5 day-old seedlings were transferred to a culture chamber (Lab-Tek, Chamberes #1.0 Borosilicate Coverglass System, catalog number: 155361) filled with – Fe or +Fe medium described in Gruber et al., 2013(Gruber et al., 2013) for 4 hours. After 4 hours, the cell wall was counter-stained by placing one drop of propidium iodide 15 µM (10 µg/mL in distilled water) on the root tip for 1 minutes. A coverslip was placed on the surface of the root for further imaging. DRONPA-s was bleached using the full laser power (150 W) of the 488nm laser for 10 seconds. Then 2-4 regions of interest (ROIs) were drawn on the external lateral side of the epidermal root cells and DRONPA-s was activated in this region using the 405 nm laser doing 8 cycles at 15W using the FRAP module available on the Zeiss LSM 980 Airyscan 2. Right after activation and then again 6 minutes later, images were acquired in both channel, PI and DRONPA-s.

#### IRT1 reporter lines after 24h

About 24 of 5-day-old seedlings grown on iron sufficient medium were transferred to agar plate filled with +Fe or – Fe supplemented with 100 Ferrozine for 24 hours. 15 seedlings were transferred to a culture chamber (Lab-Tek, Chamberes #1.0 Borosilicate Coverglass System, catalog number: 155361) filled with +Fe described in Gruber et al., 2013(Gruber et al., 2013). The cell wall was counter-stained by placing one drop of propidium iodide 15 µM (10 µg/mL in distilled water) on the root tip for 1 minutes. A coverslip was placed on the surface of the root for further imaging. Images were acquired using the spinning disc set up described above using stitching and z-stack modes.

#### Time lapse imaging of IRT1 reporter lines

5-day-old seedlings were grown on iron sufficient medium and then about 15 seedlings were transferred to a culture chamber (Lab-Tek, Chamberes #1.0 Borosilicate Coverglass System, catalog number: 155361) filled with +Fe or – Fe medium described in Gruber et al., 2013(Gruber et al., 2013) or +Fe medium containing flg22. Note that the transfer took about 45-60 seconds. Images were acquired every 20 minutes for 16 hours representing 80 images per root using a Keyence microscope model BZ-X810 with BZ NIKON Objective Lens (2X) CFI Plan-Apo Lambda in brightfield, green (ET470/40x ET525/50m T495lpxr-UF1) or red (ET560/40x ET630/75m T585lpxr-UF1) channels.

#### Time lapse imaging of SRF3 transcriptional and translational reporter and control lines

About 15 5-day-old seedlings grown on iron sufficient medium were transferred to a culture chamber (Lab-Tek, Chamberes #1.0 Borosilicate Coverglass System, catalog number: 155361) filled with +Fe or – Fe medium described in Gruber et al., 2013. Note that the transfer took about 45-60 seconds. Images were acquired every 10 minutes for 4 hours using the spinning disc set up described above and assembled using the stitching mode, z-satck and definite focus options to keep track of the root and be localized at the same z-stage a long time, respectively.

#### Cryofixation and freeze-substitution

5-day-old seedlings of pSRF3::SRF3-GFP line (*Landsberg erecta* background) and LER were grown vertically on Caisson media complemented in iron. The seedlings were incubated for 4 hours in liquid Caisson media which were complemented or deficient in iron. Root tips were taken and cryofixed in 20% BSA filled copper platelets (100 nm deep and 1.5 mm wide) with EM PACT1 high pressure freezer (Leica). The samples were transferred for freeze-substitution in AFS2 (Leica) at −90°C in cryosubstitution mix: uranyl acetate 0.36%, in pure acetone, for 24 hours. The temperature was raised stepwise by 3°C h^-1^ until reaching −50°C and maintained for 3 hours. The cryosubstitution mix was removed and replaced by pure acetone and then pure ethanol, for each of them 3 washes of 10 minutes were performed. The copper platelets were not removed in order to avoid sample loss. HM20 Lowicryl resin (Electron Microscopy Science) solutions of increasing concentrations were used for infiltration: 25% and 50% (1 hour each), 75% (2 hours), 100% (overnight, 4 hours, 48 hours- each bath was performed with new resin). The samples were then polymerized under ultraviolet light for 24 hours at −50°C before raising the temperature stepwise by 3°C h^-1^ until reaching 20°C and maintained for 6 hours.

#### Immunogold labelling

The samples were recovered by removing exceeding resin on the top and edges of the copper platelets. The latter were removed by applying alternatively heat shocks with liquid nitrogen and on a 40°C heated knife to dissociate copper platelet from the resin. Ultrathin sections of 90 nm thickness were trimmed at a speed of 1 mm s^-1^ (EM UC7 ultramicrotome, Leica) and recovered on electron microscopy grids (T 300mesh cupper grids, Electron Microscopy Science) covered by 2% parlodion film. Once the grids were dry immunogold labelling was performed. The grids were successively incubated in 10 µl droplets of different reagents (0.22 µm filtered). The grids were first incubated in PHEM Tween 0.2% BSA 1% buffer (pH6.9) for 1 minute of rinsing before 30 minutes of blocking. The primary antibody anti-GFP rabbit polyclonal antibody (A11122, Thermo Fisher Scientific) and secondary antibody 10 nm colloidal gold-conjugated goat anti-rabbit IgG (Tebu-Bio) were diluted in PHEM Tween 0.2% BSA 1% buffer (pH6.9) to 1/200 and 1/40 respectively and grids were incubated for 1 hour. 3 rinsing steps of 5 minutes each were performed between the primary and secondary antibody incubation and after the secondary incubation. The grids were rinsed on filtered miliQ water droplets before drying and imaging. Image acquisition was performed at 42000x magnification on a FEI Tecnai G2 Spirit TWIN TEM with axial Eagle 4K camera.

#### Immunolocalization of callose

Arabidopsis seedlings were grown on ½ MS 1% sucrose agar plate for 6 days and then incubated for 3 hours in ½ MS 1% sucrose liquid medium for control condition or ½ MS 1% sucrose liquid medium containing 0.4 M mannitol, prior to fixation. The immunolocalization procedure was done according to Boutté *et al*. 2014 (Boutté and Grebe, 2014). The callose antibody (Australia Biosupplies) was diluted to 1/300 in MTSB (Microtubule Stabilizing Buffer) containing 5% of neutral donkey serum. The secondary anti-mouse antibody coupled to TRITC (tetramethylrhodamine) was diluted to 1/300 in MTSB buffer containing 5% of neutral donkey serum. The nuclei were stained using DAPI (4’,6-diamidino-2-phénylindole) diluted to 1/200 in MTSB buffer for 20 minutes. Samples were then imaged with a Zeiss LSM 880 using X40 oil lens. DAPI excitation was performed using 0,5% of 405 laser power and fluorescence collected at 420-480 nm; GFP excitation was performed using 5% of 488 nm laser power and fluorescence emission collected at 505-550 nm; TRITC excitation was performed with 5% of 561 nm power and fluorescence collected at 569-590 nm. All the parameters were kept the same between experiments to allow quantifications.

#### Short-term iron deficiency, flg20 and flg22 treatments

Seeds were sowed in +Fe medium described in Gruber et al., 2013 and stratified for 2-3 days at 4°C. 5-day-old seedlings were treated for 4 hours with low iron medium or for 2 hours adding 100uM of FerroZine or sufficiency in liquid medium described in Gruber et al., 2013 using 12-well plates. Note that after the addition of FerroZine the pH was adjusted to the same pH=5.7 as the control medium +Fe. However, no change in the pH was detected in agar adjusted with MES as described earlier and in Gruber et al., 2013. For flg22 treatment, Seeds were sowed in +Fe medium described in Gruber et al., 2013 and stratified for 2-3 days at 4°C. 5-day-old seedlings were treated for 4 hours in iron sufficient media described in Gruber et al., 2013 supplemented or not with flg22 or flg20.

#### RhoNox-1 staining

5-day-old seedlings were treated in ultra-pure distilled water (Fisher Scientific Invitrogen UltraPure Distilled Water 500 mL Plastic Container – 10977015) called +Fe condition in order to get rid of any iron trace in water, 50uM of FerroZine was added for 30 minutes. Then, the plants were transferred to ultra-pure distilled water supplemented with 2.5uM of RhoNox-1 for 15 minutes (stock solution 5mM; https://goryochemical.com/).

#### Perls staining and DAB/H_2_O_2_ intensification

Perls staining and DAB/H_2_O_2_ intensification was performed as described previously (Roschzttardtz et al., 2009). The embryos were dissected and isolated from dry seeds previously imbibed in distilled water for 3-4 h. The embryos were then vacuum infiltrated with Perls stain solution (equal volumes of 4% (v/v) HCl and 4% (w/v) K-ferrocyanide) for 15 min and incubated for 30 min at room temperature (Stacey et al., 2008). The DAB intensification was performed as described in Meguro et al., 2007. After washing with distillated water, the embryos were incubated in a methanol solution containing 0.01 M NaN_3_ and 0.3% (v/v) H_2_O_2_ for 1 h, and then washed with 0.1 m phosphate buffer (pH 7.4). For the intensification reaction the embryos were incubated between 10 to 30 min in a 0.1 M phosphate buffer (pH 7.4) solution containing 0.025% (w/v) DAB (Sigma), 0.005% (v/v) H_2_O_2_, and 0.005% (w/v) CoCl_2_ (intensification solution). The reaction was terminated by rinsing with distilled water.

#### GUS Histochemical Assay

Transgenic seedlings carrying *pCYP71A12:GUS* were grown on ½ MS media for 4 days and seedlings were then grown in 6 well plates containing ½ MS (+Fe or −Fe) liquid media for 16 hours. Seedlings were then treated with 1 μM Flg22 or 1 μM Flg20 for 16 hours. After treatment with peptides plants were washed with 50 mM sodium phosphate buffer, pH 7. One milliliter of GUS substrate solution (50 mM sodium phosphate, pH 7, 10 mM EDTA, 0.5 mM K4[Fe(CN)6], 0.5 mM K3[Fe(CN)6], 0.5 mM X-Gluc, and 0.01% Silwet L-77) was poured in each well. The plants were vacuum infiltrated for 5 min and then incubated at 37°C for 4 h. Tissues were observed using a Discovery V8 microscope (Zeiss). Quantification of GUS signal in root tips of the stained seedlings was done using Fiji.

#### Aniline blue staining

5 day-old seedlings were incubated for 2h in iron deficient or sufficient medium(Gruber et al., 2013) with or without DDG and then transferred for 2 hours to 150 mM K_2_HPO_4_ and 0.01% aniline blue in 12-well plates wrapped in aluminum foil for light protection. Then imaging of the root epidermis in the elongation zone was performed.

#### Sterol treatments

For inhibitor experiments, 5-day-old seedlings were transferred to MS agar plates containing 50 ug/mL Fenpropimorph (https://www.caymanchem.com/; stock solution 50 ug/uL in DMSO) or 1 uM Lovastin (https://www.tocris.com/products/lovastatin_1530; stock solution 1 mM in DMSO) for 24 hours.

#### 2-deoxy-d-glucose (DDG) treatment

Seedlings were grown on iron sufficient medium and after 5 days transferred to iron sufficient or low iron medium or flg22 containing medium with or without DDG (diluted in H2O, stock 50mM used at 50uM; https://www.tocris.com/products/2-deoxy-d-glucose_4515).

### QUANTIFICATION

#### Late root growth response

Plates containing seedlings were scanned from days 5 to 9 after transfer to different media in order to acquire images for further quantification of the root growth rate per conditions. Plates were scanned using BRAT software(Slovak et al., 2014) each day and were stacked together using a macro in Fiji (Macro_Match_Align). We then calculated the root length for every day per genotype in each condition to evaluate the root growth rate in Fiji using the segmented line. We first calculated the mean of the root growth rate for each days 5 to 6, 6 to 7, 7 to 8, 8 to 9. These values were used to calculate the mean of root growth rate for 3 days. Then, we divided the mean of root growth rate for 3 days to a given media for each plant by the mean of root growth rate for 3 days after transfer to the control media for the entire related genotype. This ratio was used as the late root growth response to low iron levels Every experiment was repeated twice.

#### Early root growth response

Root length for each seedling was recorded for 12 hours taking a picture every 5 minutes and quantified using a Matlab script (Matlab_RootWalker). From these measurements, we ploted the root length from T0 to T12 after transfer. We obtain a curve representing the root length after transfer from which we calculated the area under the curve using the following formula “(Root length T1+Root length T2)/2*(Time T2-Time T1)”. Then, we divided the value of the area under the curve after transfer for each plant in a given condition by the area under the curve after transfer to the control media for the entire related genotype. This ratio was used as the early root growth response to low iron levels. Every experiment has been repeated three times.

#### Lateral root density

12-day-old seedlings were used for quantification for the lateral root assays. Plates were scanned using BRAT software(Slovak et al., 2014). A ratio of the number of lateral roots divided by the root length was applied in order to calculate the lateral root density. This final value was used for further analysis. This experiment has been repeated twice.

#### Root meristem size

5 days old seedlings were transferred to iron sufficient or deficient medium (as described in Gruber et al., 2013) that was contained in small chambers used for the early root growth response (Lab-Tek, Chamberes #1.0 Borosilicate Coverglass System, catalog number: 155361). After 12 hours, the cell wall was stained by placing one drop of propidium iodide 15 µM (10 µg/mL in distilled water) on the root tip for 5 minutes. Images were acquired using the stitching mode on the microscope. The cell size was determined using the Cell-O-Type software in the cortex cells(French et al., 2012). Every experiment has been repeated three times.

#### Measuring signal intensities at the plasma membrane

Confocal images were first denoised using an auto local threshold applying the Otsu method with a radius of 25 and a median filter with a radius of 2 in Fiji(Schindelin et al., 2012). In order to remove every single bright pixel on the generated-binary image the despeckle function was applied. In order to obtain plasma membrane skeleton, we detected and removed every intracellular dot using the “Analyze Particles” plugin with the following parameter, size between 0.0001 and 35 000 μm^2^ and a circularity between 0.18 and 1. Then, we selected and cropped a zone which only showed a proper plasma membrane skeleton. We created a selection from the generated-plasma membrane skeleton and transposed it to the original image to calculate the plasma membrane intensity. This process has been automatized in a Macro (Macro_PM_Intensity). The plasma membrane intensity value was then divided by the total intensity of the image to normalize the plasma membrane intensity. An average of 45 cells were used for quantification per root. Every experiment was repeated three times.

#### Calculating standard deviation measures of the intensities at the plasma membrane

The standard deviation of the apical-basal plasma membrane was calculated using the segmented lines in Fiji toolbox with a width of 3 pixels. 5 plasma membranes were used per root and the mean was calculated per root. Every experiment has been repeated three times.

#### Fluorescence intensity in the root tip during time lapse experiments

To acquire images, z-stacks with a stepsize of 50 μm were performed coupled with the stitching mode. To determine the variation of our translational and transcriptional reporters under different condition, we measured the signal intensity in the root tip over time using Fiji. Prior analysis, confocal images were stitched, and we generate the maximum intensity projection. We drew a region of interest of the same size in the x and y dimensions, corresponding to the width of the root and in length corresponding to the basal meristem, transition and elongation zones. In this region, for each time point we determined the mean grey value. Note that this value is normalized since for each root the same area has been kept between conditions and genotypes. Every experiment was repeated three times.

#### Polarity Index

5 days old seedlings of transgenic lines were analyzed to determine the “Polarity index” in root tip epidermis. “Polarity index” is the ratio between the fluorescence intensity (Mean Grey Value function of Fiji software) measured at the PM apical/basal side and PM lateral sides (Line width=3). We selected only cells for which the PM at each pole (apical, basal and laterals) were easily viewable and we selected cells that were entering elongation (at least as long as wide, but no more than twice as long as wide). Quantification was conducted in 100 cells over more than 15 independent plants. This Polarity index reveals the degree of polarity of the fluorescent reporters between the apical/basal side and lateral sides of the PM. Every experiment was repeated three times

#### Integrated Nuclear and fluorescence signal density of transcriptional reporter lines

To acquire images, z-stacks with a stepsize of 50 μm were performed coupled with the stitching mode. Then, we generated the maximum intensity projection for the z-dimension and then binarized the images using the auto local threshold Bernsen with a radius of 15. The despeckle and erode functions were subsequently used to remove background artefacts. The nucleus in this region were selected using the analyze particles function with the settings for size of 15 and 700 μm^2^ and for circularity 0.25 to 1.00. The regions selected were used on the original picture for determining the fluorescence intensity in each nucleus. Then the average nuclear integrated density was calculated per root in order to normalize the intensity by the total root area. This process has been automatized in a macro in Fiji (Macro_Nuclear_Signal_Intensity). From the same images, the number of nuclei was calculated and the root area was detected using the plugin “Wavelet a trou” (http://www.ens-lyon.fr/RDP/SiCE/METHODS.html) (Bayle et al., 2017). The number of nuclei detected was then divided by the area of the respective root to determine the nuclear density per root. This process was automatized using a Fiji macro (Macro_Nuclear_Density). Every experiment was repeated three times.

#### IRT1 promotor activation in time lapse serie

At each time point, roots showing fluorescence signal in the apical 3 mm of the root were counted. Roots that were already showing activation in this zone or showed slight signals at timepoint 0 were removed from the analysis. Then the number of roots that started to show a fluorescent signal after timepoint 0, was divided by the number of total roots observed in this experiment and multiplied by 100 to obtain the percentage of root showing pIRT1 activation in the apical 3 mm of the root. Every experiment was repeated three times.

#### Distance of signals from the Quiescent Center in the SRF3 transcriptional reporter line

Based on the propidium iodide staining, the Quiescent Center (QC) region was determined by its morphology. Then, using the straight-line option in Fiji, a line was traced from the QC to the first appearance of a clear, bright signal that reported pSRF3 activity. The distance along this line was calculated to determine the distance from the QC to the first cell expressing *pSRF3* in the elongation zone. Every experiment was repeated three times.

#### DRONPA-s diffusion

After activation of the DRONPA-s reporter, the signal intensity using the integrated density of the signal was calculated in the activated cells as well as the adjacent upper and lower cells using Fiji. The signal intensity once again was calculated in the same regions 6 minutes after the activation. To account for photo bleaching, these values were normalized by dividing the DRONPA-s signal intensity after and post bleach in a zone where DRONPA-s was visible in both images. The average of the normalized integrated density in the surrounding cells was calculated, averaging the values obtained in upper and lower cells. Finally, the ratio of the normalized integrated density after and post bleach was calculated by dividing both values to obtain the DRONPA-s normalized fluorescence intensity. Every experiment was repeated three times.

#### pSUC2::GFP diffusion

In order to evaluate the diffusion of the GFP protein, a ROI of about 1800 μm^2^ was drawn at the root tip of each root using Fiji. The mean gray value was calculated and divided by the corresponding area to normalize the value. Every experiment was repeated three times.

#### RhoNOx-1 signal intensity

The root area was detected using the plugin “Wavelet a trou” (http://www.ens-lyon.fr/RDP/SiCE/METHODS.html)(Bayle et al., 2017) and then in this area the mean gray value was calculated and divided by the size of the size of the area to obtain the normalized signal intensity. This process has been automatize on a Fiji Macro, Macro_RhoNox-1

#### Aniline blue fluorescence intensity

Using Fiji, the mean gray value of 10 plasma membranes in the apical-basal side of the epidermis in the transition-elongation zone was calculated with the segmented line option with 3 pixels wide. The mean of these values was then divided by the mean gray value in the total area where the plasma membrane signal has been calculated in order to normalize the value due to differential strength of the staining. This final value was used for further analysis. Every experiment was repeated three times.

#### Anti-callose antibody fluorescence intensity

Callose deposition was quantified using the Fiji software. Callose fluorescence intensity was measured at the apico-basal cell walls of root epidermal and cortex cells using the segmented line with a width of 3 pixels. A total of 8 to 10 cell walls were measured per roots and used to calculate the average of anti-callose fluorescence intensity per root. Between 5 to 20 roots per transgenic lines and conditions were used in two independent biological replicates were used.

#### Immunogold

The number of gold particles in the EM micrographs were quantified using Fiji software. The number of particles were counted manually in each compartment; e.g. Cytosol, Cell wall, PM and PD; and then reported relative to the surface of cytosol or cell wall (/ m^2^), to the length of PM (/ m) and to individual PD. A total of 25 to 50 micrographs were analyzed for each line and conditions. Two biological replicates were used.

### STATISTICS

Each experiment has been repeated independently at least twice, as in every cases the same trend has been recorded for independent experiment, the data the different has been pooled for further statistical analysis. Each sample were subjected to four different normality tests (Jarque-Bera, Lilliefors, Anderson-Darling and Shapiro-Wilk), sample were considered as a Gaussian distribution when at least one test was significant (p=0.05) using Xlstat.

- As a normal distribution was observed a one-way ANOVA coupled with post hoc Tukey honestly significant difference (HSD) test was performed (p=0.05) using R software or Xlstat. Figures: 1F, 2A, 3A, 4A, 4B, 4C, 4D, 5E, 6B, S2C, S2D, S2F, S3C, S4A, S4B, S8C, S9D, S11A, S11B, S12B, S12C, S12D, S12E, S13D, S13E, S17C, S17D.
- As a normal distribution was observed at one-way ANOVA coupled with post hoc Lowest significant difference (LSD) test was performed (p=0.05) using Xlstat: 5E.
- As a normal distribution was observed an independent two-ways student test was performed (p=0.05) using Xlstat. Figures: 1C, 1D, 2D, 5D, 6C, 6D, S2E, S4C, S7B (left panel), S7C, S10A, S10B, S10C, S11C, S14A, S14C (both panel).
- As a normal distribution was not observed at two-ways Kruskal-Wallis coupled with post hoc Steel-Dwass-Critchlow-Fligner procedure was performed (p=0.05) using Xlstat. Figure: S9C.
- As a normal distribution was not observed a two-ways Mann-Whitney test was performed (p=0.05) using Xlstat. Figures: S7B (right panel)

For time lapse analysis SAS software was used based on a mixed effect model (*p<0.05*) to test the statistical significance. Figures: 2C

### CLONING

#### pIRT1 transcriptional reporter line

To generate the transcriptional pIRT1::NLS-2xYPet reporter line, the *IRT1* promoter (2.6 kb) was cloned at *Sal*I and *Bam*HI restriction sites using the primers pIRT1 Sal_F and pIRT1 Bam_R in the pBJ36 vector carrying two in frame copies of the YPet yellow fluorescent protein fused to SV40 nuclear localization signal (kind gift of Dr. Jeff D.B. Long, UCLA). The *pIRT1::2xYPet-NLS* cassette was digested with *Not*I and cloned in the pART27 binary vector(Gleave, 1992). Note that About 20 independent T1 lines were isolated and between three to six representative mono-insertion lines showing strong activation of *IRT1* promoter in the root epidermis upon low iron, as previously described (Vert et al., 2002), were selected in T2.

#### SRF3 constructs in Ler

To generate pSRF3::SRF3g-GFP from *Ler* background, the SRF3 gene and its native promoter (−1492 nt from the transcription start) was amplified by PCR using genomic DNA as template and the primers attB1-SRF3Ler_F and attB2-SRF3Ler_R. The resulting amplicon was purified, sequenced and subcloned into pDONR221 by Gateway BP recombination, following manufacturer’s instructions. To generate the C-terminus GFP fusion, the pSRF3::SRF3g fragment was cloned into the binary vector pGWB450 (Nakagawa et al., 2007) by Gateway LR recombination.

#### SRF3 constructs (entry vectors)

The full-length coding sequence of SRF3 (At4g03390) was amplified by RT-PCR using 7-day old Arabidopsis seedlings cDNA as template and the SRF3_CDS_p221_F and PSRF3_CDS_p221_noSTOP_R primers. The corresponding PCR product was recombined into pDONR221 vector by BP reaction to give SRF3cds-noSTOP/pDONR221. To remove the SRF3 extracellular domain the primers SRF3_kinase_p221_F and SRF3_kinase_p221_R. The corresponding PCR product was recombined into pDONR221 vector by BP reaction to give SRF3cdsΔextraC_p221.

To remove the SRF3 kinase domain 5’ phosphorylated primers were used SRF3_ΔKinase2-5’_F and SRF3_ΔKinase-5’_R followed by a ligation to give SRF3cdsΔKinase_pDONR221.

SRF3 mutant impaired in the kinase activity was obtained by site directed mutagenesis using SRF3-cds_mutKD_p221_F and SRF3-cds_mutKD_p221_R to give SRF3cds KDmut_pDONR221.

#### Promoters and fluorescent proteins (entry vectors)

The *SRF3* promoter (5078bp upstream of 5’UTR until the 3’UTR of the previous gene) was cloned using the gibson cloning method (https://www.neb.com/applications/cloning-and-synthetic-biology/dna-assembly-and-cloning/gibson-assembly#tabselect3) with the following primers, Insert_pSRF3_F, Insert_pSRF3_R and Backbone_pSRF3_F and Backbone_pSRF3_R introduced into the P4P1R vector (life technologies www.lifetechnologies.com/) to give SRF3prom/pDONR4P1R.

The fluorescent mSCARLET protein was synthesized (GeneArt, www.thermofischer.com), amplified with attB2r and attB3 gateway sites using the mSCARLET_F and mSCARLETwSTOP_R primers, and then recombined into pDONRP2R-P3 by BP reaction to yield the mSCARLET/pDONRP2R-P3 entry vector.

## References

Alcázar, R., García, A.V., Kronholm, I., Meaux, J. de, Koornneef, M., Parker, J.E., and Reymond, M. (2010). Natural variation at Strubbelig Receptor Kinase 3 drives immune-triggered incompatibilities between *Arabidopsis thaliana* accessions. Nature Genetics 42, 1135–1139.

Anders, S., Pyl, P.T., and Huber, W. (2014). HTSeq—a Python framework to work with high-throughput sequencing data. Bioinformatics 31, 166–169.

Atwell, S., Huang, Y.S., Vilhjálmsson, B.J., Willems, G., Horton, M., Li, Y., Meng, D., Platt, A., Tarone, A.M., Hu, T.T., et al. (2010). Genome-wide association study of 107 phenotypes in Arabidopsis thaliana inbred lines. Nature 465, 627–631.

Aznar, A., Chen, N.W.G., Rigault, M., Riache, N., Joseph, D., Desmaële, D., Mouille, G., Boutet, S., Soubigou-Taconnat, L., Renou, J.-P., et al. (2014). Scavenging Iron: A Novel Mechanism of Plant Immunity Activation by Microbial Siderophores1[C][W]. Plant Physiol 164, 2167–2183.

Bai, L., Qiao, M., Zheng, R., Deng, C., Mei, S., and Chen, W. (2016). Phylogenomic analysis of transferrin family from animals and plants. Comp Biochem Physiol Part D Genomics Proteomics 17, 1–8.

Bayle, V., Platre, M.P., and Jaillais, Y. (2017). Automatic Quantification of the Number of Intracellular Compartments in Arabidopsis thaliana Root Cells. BIO-PROTOCOL 7.

Belkhadir, Y., and Jaillais, Y. (2015). The molecular circuitry of brassinosteroid signaling. New Phytologist 206, 522–540.

Benitez-Alfonso, Y., Faulkner, C., Pendle, A., Miyashima, S., Helariutta, Y., and Maule, A. (2013). Symplastic Intercellular Connectivity Regulates Lateral Root Patterning. Developmental Cell 26, 136–147.

Benjamini, Y., and Yekutieli, D. (2001). The control of the false discovery rate in multiple testing under dependency. Ann. Statist. 29, 1165–1188.

Benschop, J.J., Mohammed, S., and Slijper, M. (2007). Quantitative Phosphoproteomics of Early Elicitor Signaling in Arabidopsis. Mol Cell Proteomics 17.

Boutté, Y., and Grebe, M. (2014). Immunocytochemical fluorescent in situ visualization of proteins in arabidopsis. Methods in Molecular Biology 1062, 453–472.

Brachi, B., Faure, N., Horton, M., Flahauw, E., Vazquez, A., Nordborg, M., Bergelson, J., Cuguen, J., and Roux, F. (2010). Linkage and Association Mapping of Arabidopsis thaliana Flowering Time in Nature. PLOS Genetics 6, e1000940.

Cassat, J.E., and Skaar, E.P. (2013). Iron in Infection and Immunity. Cell Host & Microbe 13, 509–519.

Cheval, C., Samwald, S., Johnston, M.G., Keijzer, J. de, Breakspear, A., Liu, X., Bellandi, A., Kadota, Y., Zipfel, C., and Faulkner, C. (2020). Chitin perception in plasmodesmata characterizes submembrane immune-signaling specificity in plants. PNAS 117, 9621–9629.

Clough, S.J., and Bent, A.F. (1998). Floral dip: a simplified method for Agrobacterium-mediated transformation of Arabidopsis thaliana. Plant J 16, 735–743.

Cutler, S.R., Ehrhardt, D.W., Griffitts, J.S., and Somerville, C.R. (2000). Random GFP∷cDNA fusions enable visualization of subcellular structures in cells of Arabidopsis at a high frequency. PNAS 97, 3718–3723.

Deák, M., Horváth, G.V., Davletova, S., Török, K., Sass, L., Vass, I., Barna, B., Király, Z., and Dudits, D. (1999). Plants ectopically expressing the iron-binding protein, ferritin, are tolerant to oxidative damage and pathogens. Nat. Biotechnol. 17, 192–196.

Denyer, T., Xiaoli, M., Klesen, S., Scacchi, E., Nieselt, K., and Timmermans, M.C.P. (2019). Spatiotemporal Developmental Trajectories in the Arabidopsis Root Revealed Using High-Throughput Single-Cell RNA Sequencing. Developmental Cell 49, 19.

Dinneny, J.R., Long, T.A., Wang, J.Y., Jung, J.W., Mace, D., Pointer, S., Barron, C., Brady, S.M., Schiefelbein, J., and Benfey, P.N. (2008). Cell identity mediates the response of Arabidopsis roots to abiotic stress. Science 320, 942–945.

Durrett, T.P., Gassmann, W., and Rogers, E.E. (2007). The FRD3-mediated efflux of citrate into the root vasculature is necessary for efficient iron translocation. Plant Physiol 144, 197–205.

Fichman, Y., Myers, R.J., Grant, D.G., and Mittler, R. (2021). Plasmodesmata-localized proteins and ROS orchestrate light-induced rapid systemic signaling in Arabidopsis. Sci. Signal. 14.

French, A.P., Wilson, M.H., Kenobi, K., Dietrich, D., Voss, U., Ubeda-Tomas, S., Pridmore, T.P., and Wells, D.M. (2012). Identifying biological landmarks using a novel cell measuring image analysis tool: Cell-o-Tape. Plant Methods 8, 7.

Ganz, T., and Nemeth, E. (2015). Iron homeostasis in host defence and inflammation. Nat Rev Immunol 15, 500–510.

García, M.J., Romera, F.J., Stacey, M.G., Stacey, G., Villar, E., Alcántara, E., and Pérez-Vicente, R. (2013). Shoot to root communication is necessary to control the expression of iron-acquisition genes in Strategy I plants. Planta 237, 65–75.

Gerlitz, N., Gerum, R., Sauer, N., and Stadler, R. (2018). Photoinducible DRONPA-s: a new tool for investigating cell-cell connectivity. Plant J 94, 751–766.

Gleave, A.P. (1992). A versatile binary vector system with a T-DNA organisational structure conducive to efficient integration of cloned DNA into the plant genome. Plant Mol Biol 20, 1203–1207.

Grillet, L., Lan, P., Li, W., Mokkapati, G., and Schmidt, W. (2018). IRON MAN is a ubiquitous family of peptides that control iron transport in plants. Nature Plants 4, 953–963.

Grison, M.S., Brocard, L., Fouillen, L., Nicolas, W., Wewer, V., Dörmann, P., Nacir, H., Benitez-Alfonso, Y., Claverol, S., Germain, V., et al. (2015). Specific Membrane Lipid Composition Is Important for Plasmodesmata Function in Arabidopsis. The Plant Cell 27, 1228–1250.

Grison, M.S., Kirk, P., Brault, M.L., Wu, X.N., Schulze, W.X., Benitez-Alfonso, Y., Immel, F., and Bayer, E.M. (2019). Plasma Membrane-Associated Receptor-like Kinases Relocalize to Plasmodesmata in Response to Osmotic Stress. Plant Physiology 181, 142–160.

Groen, S.C., Whiteman, N.K., Bahrami, A.K., Wilczek, A.M., Cui, J., Russell, J.A., Cibrian-Jaramillo, A., Butler, I.A., Rana, J.D., Huang, G.-H., et al. (2013). Pathogen-Triggered Ethylene Signaling Mediates Systemic-Induced Susceptibility to Herbivory in Arabidopsis[W]. Plant Cell 25, 4755–4766.

Gruber, B.D., Giehl, R.F.H., Friedel, S., and von Wirén, N. (2013). Plasticity of the Arabidopsis Root System under Nutrient Deficiencies. Plant Physiol. 163, 161–179.

Han, X., Hyun, T.K., Zhang, M., Kumar, R., Koh, E., Kang, B.-H., Lucas, W.J., and Kim, J.-Y. (2014). Auxin-Callose-Mediated Plasmodesmal Gating Is Essential for Tropic Auxin Gradient Formation and Signaling. Developmental Cell 28, 132–146.

Haydon, M.J., Kawachi, M., Wirtz, M., Hillmer, S., Hell, R., and Krämer, U. (2012). Vacuolar nicotianamine has critical and distinct roles under iron deficiency and for zinc sequestration in Arabidopsis. Plant Cell 24, 724–737.

He, P., Shan, L., Lin, N.-C., Martin, G.B., Kemmerling, B., Nürnberger, T., and Sheen, J. (2006). Specific Bacterial Suppressors of MAMP Signaling Upstream of MAPKKK in Arabidopsis Innate Immunity. Cell 125, 563–575.

Higashi, K., Ishiga, Y., Inagaki, Y., Toyoda, K., Shiraishi, T., and Ichinose, Y. (2008). Modulation of defense signal transduction by flagellin-induced WRKY41 transcription factor in Arabidopsis thaliana. Mol Genet Genomics 279, 303–312.

Hindt, M.N., Akmakjian, G.Z., Pivarski, K.L., Punshon, T., Baxter, I., Salt, D.E., and Guerinot, M.L. (2017). BRUTUS and its paralogs, BTS LIKE1 and BTS LIKE2, encode important negative regulators of the iron deficiency response in Arabidopsis thaliana. Metallomics 9, 876–890.

Hirayama, T. (2018). Development of Chemical Tools for Imaging of Fe(II) Ions in Living Cells: A Review. Acta Histochem Cytochem 51, 137–143.

Hohmann, U., Lau, K., and Hothorn, M. (2017). The Structural Basis of Ligand Perception and Signal Activation by Receptor Kinases (Annual Reviews).

Horton, M.W., Hancock, A.M., Huang, Y.S., Toomajian, C., Atwell, S., Auton, A., Muliyati, N.W., Platt, A., Sperone, F.G., Vilhjálmsson, B.J., et al. (2012). Genome-wide patterns of genetic variation in worldwide Arabidopsis thaliana accessions from the RegMap panel. Nature Genetics 44, 212–216.

Huang, D., Sun, Y., Ma, Z., Ke, M., Cui, Y., Chen, Z., Chen, C., Ji, C., Tran, T.M., Yang, L., et al. (2019). Salicylic acid-mediated plasmodesmal closure via Remorin-dependent lipid organization. PNAS 116, 21274–21284.

Iatsenko, I., Marra, A., Boquete, J.-P., Peña, J., and Lemaitre, B. (2020). Iron sequestration by transferrin 1 mediates nutritional immunity in Drosophila melanogaster. PNAS 117, 7317–7325.

Jaffe, M.J., and Leopold, A.C. (1984). Callose deposition during gravitropism of Zea mays and Pisum sativum and its inhibition by 2-deoxy-D-glucose. Planta 161, 20–26.

Jaillais, Y., and Vert, G. (2016). Brassinosteroid signaling and BRI1 dynamics went underground. Current Opinion in Plant Biology 33, 92–100.

Jaillais, Y., Hothorn, M., Belkhadir, Y., Dabi, T., Nimchuk, Z.L., Meyerowitz, E.M., and Chory, J. (2011). Tyrosine phosphorylation controls brassinosteroid receptor activation by triggering membrane release of its kinase inhibitor. Genes Dev. 25, 232–237.

Kang, H.M., Zaitlen, N.A., Wade, C.M., Kirby, A., Heckerman, D., Daly, M.J., and Eskin, E. (2008). Efficient Control of Population Structure in Model Organism Association Mapping. Genetics 178, 1709–1723.

Karimi, M., Depicker, A., and Hilson, P. (2007). Recombinational Cloning with Plant Gateway Vectors. Plant Physiology 145, 1144–1154.

Khan, M.A., Castro-Guerrero, N.A., McInturf, S.A., Nguyen, N.T., Dame, A.N., Wang, J., Bindbeutel, R.K., Joshi, T., Jurisson, S.S., Nusinow, D.A., et al. (2018). Changes in iron availability in Arabidopsis are rapidly sensed in the leaf vasculature and impaired sensing leads to opposite transcriptional programs in leaves and roots. Plant Cell Environ 41, 2263–2276.

Kim, D., Pertea, G., Trapnell, C., Pimentel, H., Kelley, R., and Salzberg, S.L. (2013). TopHat2: accurate alignment of transcriptomes in the presence of insertions, deletions and gene fusions. Genome Biology 14, R36.

Kim, S.A., Punshon, T., Lanzirotti, A., Li, L., Alonso, J.M., Ecker, J.R., Kaplan, J., and Guerinot, M.L. (2006). Localization of iron in Arabidopsis seed requires the vacuolar membrane transporter VIT1. Science 314, 1295–1298.

Klatte, M., Schuler, M., Wirtz, M., Fink-Straube, C., Hell, R., and Bauer, P. (2009). The Analysis of Arabidopsis Nicotianamine Synthase Mutants Reveals Functions for Nicotianamine in Seed Iron Loading and Iron Deficiency Responses. Plant Physiology 150, 257.

Kobayashi, T., and Nishizawa, N.K. (2012). Iron Uptake, Translocation, and Regulation in Higher Plants. Annual Review of Plant Biology 63, 131–152.

Koen, E., Besson-Bard, A., Duc, C., Astier, J., Gravot, A., Richaud, P., Lamotte, O., Boucherez, J., Gaymard, F., and Wendehenne, D. (2013). Arabidopsis thaliana nicotianamine synthase 4 is required for proper response to iron deficiency and to cadmium exposure. Plant Sci 209, 1–11.

Kramer, J., Özkaya, Ö., and Kümmerli, R. (2020). Bacterial siderophores in community and host interactions. Nature Reviews Microbiology 18, 152–163.

Kreps, J.A., Wu, Y., Chang, H.-S., Zhu, T., Wang, X., and Harper, J.F. (2002). Transcriptome Changes for Arabidopsis in Response to Salt, Osmotic, and Cold Stress. Plant Physiol 130, 2129–2141.

Kumar, R.K., Chu, H.-H., Abundis, C., Vasques, K., Rodriguez, D.C., Chia, J.-C., Huang, R., Vatamaniuk, O.K., and Walker, E.L. (2017). Iron-Nicotianamine Transporters Are Required for Proper Long Distance Iron Signaling. Plant Physiology 175, 15.

Lee, J.-Y., Wang, X., Cui, W., Sager, R., Modla, S., Czymmek, K., Zybaliov, B., Wijk, K. van, Zhang, C., Lu, H., et al. (2011a). A Plasmodesmata-Localized Protein Mediates Crosstalk between Cell-to-Cell Communication and Innate Immunity in Arabidopsis. The Plant Cell Online 23, 3353–3373.

Lee, J.-Y., Wang, X., Cui, W., Sager, R., Modla, S., Czymmek, K., Zybaliov, B., van Wijk, K., Zhang, C., Lu, H., et al. (2011b). A Plasmodesmata-Localized Protein Mediates Crosstalk between Cell-to-Cell Communication and Innate Immunity in Arabidopsis. The Plant Cell 23, 22.

Lim, G.-H., Shine, M.B., de Lorenzo, L., Yu, K., Cui, W., Navarre, D., Hunt, A.G., Lee, J.-Y., Kachroo, A., and Kachroo, P. (2016). Plasmodesmata Localizing Proteins Regulate Transport and Signaling during Systemic Acquired Immunity in Plants. Cell Host & Microbe 19, 541–549.

Marquès-Bueno, M.M., Morao, A.K., Cayrel, A., Platre, M.P., Barberon, M., Caillieux, E., Colot, V., Jaillais, Y., Roudier, F., and Vert, G. (2016). A versatile Multisite Gateway-compatible promoter and transgenic line collection for cell type-specific functional genomics in Arabidopsis. The Plant Journal 85, 320–333.

Mendoza-Cózatl, D.G., Xie, Q., Akmakjian, G.Z., Jobe, T.O., Patel, A., Stacey, M.G., Song, L., Demoin, D.W., Jurisson, S.S., Stacey, G., et al. (2014). OPT3 Is a Component of the Iron-Signaling Network between Leaves and Roots and Misregulation of OPT3 Leads to an Over-Accumulation of Cadmium in Seeds. Mol Plant 7, 1455–1469.

Millet, Y.A., Danna, C.H., Clay, N.K., Songnuan, W., and Simon, M.D. (2010). Innate Immune Responses Activated in Arabidopsis Roots by Microbe-Associated Molecular Patterns W OA. The Plant Cell 22, 18.

Nemoto, K., Seto, T., Takahashi, H., Nozawa, A., Seki, M., Shinozaki, K., Endo, Y., and Sawasaki, T. (2011). Autophosphorylation profiling of Arabidopsis protein kinases using the cell-free system. Phytochemistry 72, 1136–1144.

Nicolas, W.J., Grison, M.S., Trépout, S., Gaston, A., Fouché, M., Cordelières, F.P., Oparka, K., Tilsner, J., Brocard, L., and Bayer, E.M. (2017). Architecture and permeability of post-cytokinesis plasmodesmata lacking cytoplasmic sleeves. Nat Plants 3, 17082.

Palmer, C.M., Hindt, M.N., Schmidt, H., Clemens, S., and Guerinot, M.L. (2013). MYB10 and MYB72 Are Required for Growth under Iron-Limiting Conditions. PLOS Genetics 9, e1003953.

Perraki, A., DeFalco, T.A., Derbyshire, P., Avila, J., Séré, D., Sklenar, J., Qi, X., Stransfeld, L., Schwessinger, B., Kadota, Y., et al. (2018). Phosphocode-dependent functional dichotomy of a common co-receptor in plant signalling. Nature 561, 248–252.

Platre, M.P., Noack, L.C., Doumane, M., Bayle, V., Simon, M.L.A., Maneta-Peyret, L., Fouillen, L., Stanislas, T., Armengot, L., Pejchar, P., et al. (2018). A Combinatorial Lipid Code Shapes the Electrostatic Landscape of Plant Endomembranes. Dev Cell 45, 465–480.e11.

Robinson, M.D., McCarthy, D.J., and Smyth, G.K. (2009). edgeR: a Bioconductor package for differential expression analysis of digital gene expression data. Bioinformatics 26, 139–140.

Rutschow, H.L., Baskin, T.I., and Kramer, E.M. (2011). Regulation of Solute Flux through Plasmodesmata in the Root Meristem. Plant Physiology 155, 1817–1826.

Sager, R.E., and Lee, J.-Y. (2018). Plasmodesmata at a glance. J Cell Sci 131.

Sager, R., Wang, X., Hill, K., Yoo, B.-C., Caplan, J., Nedo, A., Tran, T., Bennett, M.J., and Lee, J.-Y. (2020). Auxin-dependent control of a plasmodesmal regulator creates a negative feedback loop modulating lateral root emergence. Nature Communications 11, 1–10.

Satbhai, S.B., Setzer, C., Freynschlag, F., Slovak, R., Kerdaffrec, E., and Busch, W. (2017). Natural allelic variation of FRO2 modulates Arabidopsis root growth under iron deficiency. Nature Communications 8, 15603.

Schindelin, J., Arganda-Carreras, I., Frise, E., Kaynig, V., Longair, M., Pietzsch, T., Preibisch, S., Rueden, C., Saalfeld, S., Schmid, B., et al. (2012). Fiji: an open-source platform for biological-image analysis. Nat Methods 9, 676–682.

Segond, D., Dellagi, A., Lanquar, V., Rigault, M., Patrit, O., Thomine, S., and Expert, D. (2009). NRAMP genes function in Arabidopsis thaliana resistance to Erwinia chrysanthemi infection. The Plant Journal 58, 195–207.

Selote, D., Samira, R., Matthiadis, A., Gillikin, J.W., and Long, T.A. (2015). Iron-Binding E3 Ligase Mediates Iron Response in Plants by Targeting Basic Helix-Loop-Helix Transcription Factors. Plant Physiol. 167, 273–286.

Seren, Ü., Vilhjálmsson, B.J., Horton, M.W., Meng, D., Forai, P., Huang, Y.S., Long, Q., Segura, V., and Nordborg, M. (2012). GWAPP: A Web Application for Genome-Wide Association Mapping in Arabidopsis. The Plant Cell 24, 4793–4805.

Shikanai, Y., Yoshida, R., Hirano, T., Enomoto, Y., Li, B., Asada, M., Yamagami, M., Yamaguchi, K., Shigenobu, S., Tabata, R., et al. (2020). Callose Synthesis Suppresses Cell Death Induced by Low-Calcium Conditions in Leaves. Plant Physiology 182, 2199–2212.

Simon, M.L.A., Platre, M.P., Assil, S., van Wijk, R., Chen, W.Y., Chory, J., Dreux, M., Munnik, T., and Jaillais, Y. (2014). A multi-colour/multi-affinity marker set to visualize phosphoinositide dynamics in Arabidopsis. Plant J 77, 322–337.

Slovak, R., Goschl, C., Su, X., Shimotani, K., Shiina, T., and Busch, W. (2014). A Scalable Open-Source Pipeline for Large-Scale Root Phenotyping of Arabidopsis. The Plant Cell 26, 2390–2403.

Smakowska-Luzan, E., Mott, G.A., Parys, K., Stegmann, M., Howton, T.C., Layeghifard, M., Neuhold, J., Lehner, A., Kong, J., Grünwald, K., et al. (2018). An extracellular network of Arabidopsis leucine-rich repeat receptor kinases. Nature 553, 342–346.

Stonebloom, S., Brunkard, J.O., Cheung, A.C., Jiang, K., Feldman, L., and Zambryski, P. (2012). Redox States of Plastids and Mitochondria Differentially Regulate Intercellular Transport via Plasmodesmata. Plant Physiology 158, 190–199.

Stringlis, I.A., Proietti, S., Hickman, R., Verk, M.C.V., Zamioudis, C., and Pieterse, C.M.J. (2018). Root transcriptional dynamics induced by beneficial rhizobacteria and microbial immune elicitors reveal signatures of adaptation to mutualists. The Plant Journal 93, 166–180.

Tang, D., Wang, G., and Zhou, J.-M. (2017). Receptor Kinases in Plant-Pathogen Interactions: More Than Pattern Recognition. The Plant Cell 29, 618–637.

Thomas, C.L., Bayer, E.M., Ritzenthaler, C., Fernandez-Calvino, L., and Maule, A.J. (2008a). Specific Targeting of a Plasmodesmal Protein Affecting Cell-to-Cell Communication. PLOS Biology 6, e7.

Thomas, C.L., Bayer, E.M., Ritzenthaler, C., Fernandez-Calvino, L., and Maule, A.J. (2008b). Specific Targeting of a Plasmodesmal Protein Affecting Cell-to-Cell Communication. PLoS Biology 6.

Tian, T., Liu, Y., Yan, H., You, Q., Yi, X., Du, Z., Xu, W., and Su, Z. (2017). agriGO v2.0: a GO analysis toolkit for the agricultural community, 2017 update. Nucleic Acids Research 45, W122–W129.

Vatén, A., Dettmer, J., Wu, S., Stierhof, Y.-D., Miyashima, S., Yadav, S.R., Roberts, C.J., Campilho, A., Bulone, V., Lichtenberger, R., et al. (2011a). Callose Biosynthesis Regulates Symplastic Trafficking during Root Development. Developmental Cell 21, 1144–1155.

Vatén, A., Dettmer, J., Wu, S., Stierhof, Y.-D., Miyashima, S., Yadav, S.R., Roberts, C.J., Campilho, A., Bulone, V., Lichtenberger, R., et al. (2011b). Callose biosynthesis regulates symplastic trafficking during root development. Dev Cell 21, 1144–1155.

Verbon, E.H., Trapet, P.L., Stringlis, I.A., and Kruijs, S. (2017). Iron and Immunity. Annual Review of Phytopathology 55, 1–15.

Vert, G.A., Briat, J.-F., and Curie, C. (2003). Dual Regulation of the Arabidopsis High-Affinity Root Iron Uptake System by Local and Long-Distance Signals. Plant Physiol 132, 796–804.

Xing, Y., Xu, N., Bhandari, D.D., Lapin, D., Sun, X., Luo, X., Cao, J., Wang, H., Coaker, G., Parker, J.E., et al. (2021). Bacterial effector targeting of a plant iron sensor facilitates iron acquisition and pathogen. The Plant Cell Online 31.

Zamioudis, C., Korteland, J., Van Pelt, J.A., van Hamersveld, M., Dombrowski, N., Bai, Y., Hanson, J., Van Verk, M.C., Ling, H.-Q., Schulze-Lefert, P., et al. (2015). Rhizobacterial volatiles and photosynthesis-related signals coordinate MYB72 expression in Arabidopsis roots during onset of induced systemic resistance and iron-deficiency responses. Plant J. 84, 309–322.

