## Supplemental for "The receptor kinase SRF3 coordinates iron-level and flagellin dependent defense and growth responses in plants"

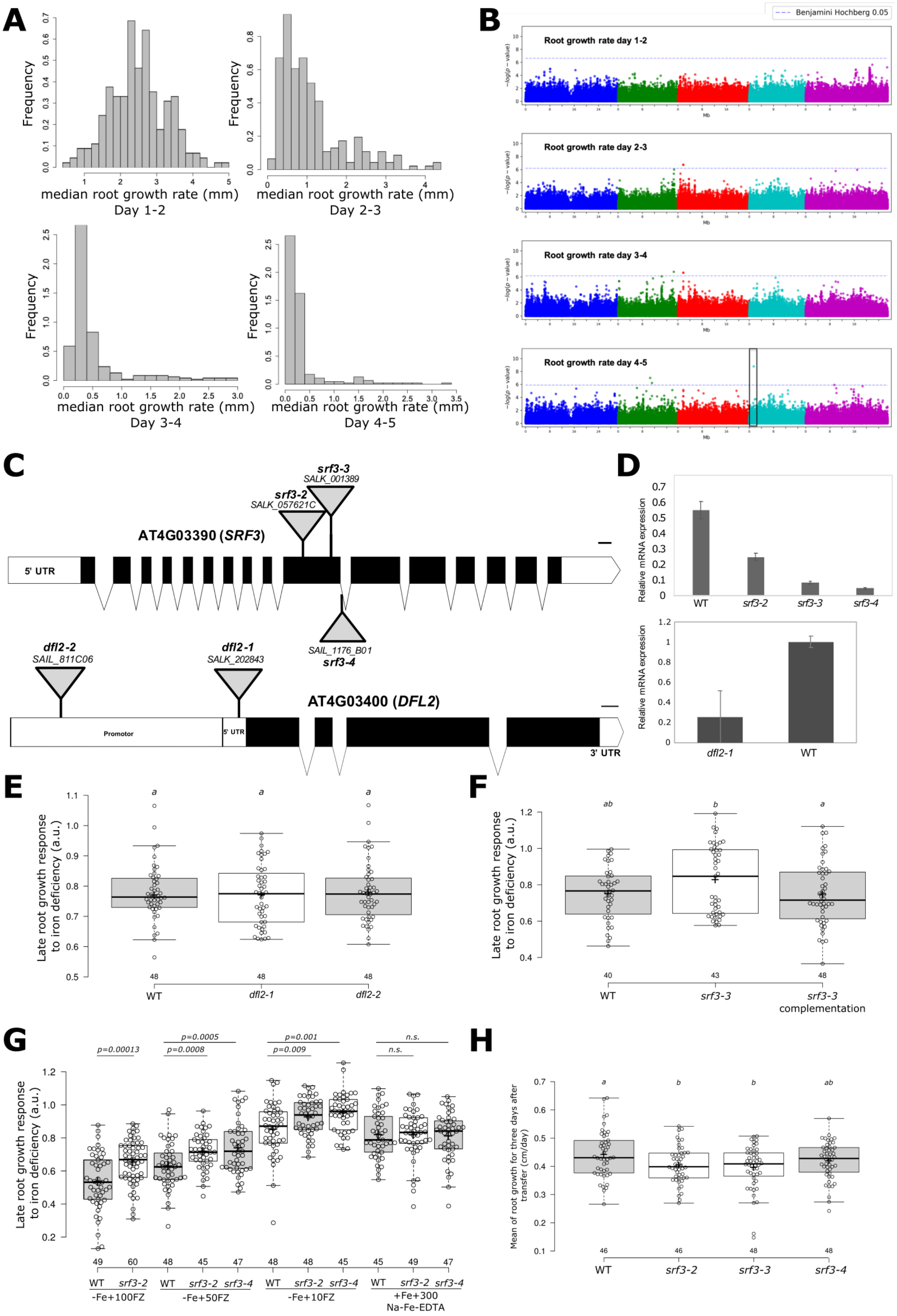
A
Frequency
median root growth rate (mm)
Day 1-2
Day 2-3
Day 3-4
Day 4-5
B
Benjamini Hochberg 0.05
Root growth rate day 1-2
Root growth rate day 2-3
Root growth rate day 3-4
Root growth rate day 4-5
−log(p − value)
Mb
C
srf3-3
SALK_001389
srf3-2
SALK_057621C
AT4G03390 (SRF3)
5' UTR
SAIL_1176_B01
srf3-4
dfl2-2
SAIL_811C06
dfl2-1
SALK_202843
AT4G03400 (DFL2)
Promotor
5' UTR
3' UTR
D
Relative mRNA expression
WT
srf3-2
srf3-3
srf3-4
dfl2-1
WT
E
Late root growth response to iron deficiency (a.u.)
a
a
a
48
48
48
WT
dfl2-1
dfl2-2
F
ab
b
a
40
43
48
WT
srf3-3
srf3-3 complementation
G
p=0.00013
p=0.0005
p=0.0008
p=0.001
p=0.009
n.s.
n.s.
49
60
48
45
47
48
48
45
45
49
47
WT
srf3-2
WT
srf3-2
srf3-4
WT
srf3-2
srf3-4
WT
srf3-2
srf3-4
-Fe+100FZ
-Fe+50FZ
-Fe+10FZ
+Fe+300 Na-Fe-EDTA
H
Mean of root growth for three days after transfer (cm/day)
a
b
b
ab
46
46
48
48
WT
srf3-2
srf3-3
srf3-4

**Supplementary Figure 1 (Related to Fig1). Root growth rate distribution GWAS for Arabidopsis accessions grown under–Fe used in this study and characterization of *srf3* mutants with srf3-neigbouring gene under low iron. (A)** Histograms of median root growth rates (mm) of accessions grown on Fe deficient growth conditions (root growth rate day 1-2, root growth rate day 2-3, root growth rate day 3-4, root growth rate day 4-5). x-axis: median root growth rate in mm; y-axis: frequency. **(B)** Manhattan plots depicting genome wide SNP associations for median root growth rate on Fe deficient growth conditions. The chromosomes are depicted in different colors. The horizontal blue dash-dot line corresponds to a nominal 0.05 significance threshold after Benjamini-Hochberg Correction. Black box indicates the significantly associated region in close proximity to *SRF3* gene. x-axis: chromosomal position of SNP; y-axis: $-\log_{10}(p\text{-value})$. **(C)** Cartoon showing the genomic structures of (a) AT4G03390 and (b) AT4G03400 and the T-DNA insertion sites**.** Black boxes indicate exons. Scale bar: 100 bp. Gene models were generated by Exon-Intron graphic maker (http://wormweb.org/exonintron). **(D)** Transcript analysis of AT4G03390 in *srf3-2, srf3-3* and *srf3-4* and transcript analysis of AT4G03400 in *at4g03400.* **(E, F)** Root growth response doing transfer assay under low iron levels supplemented with 100µM ferrozine with WT, SALK_202843 and SAIL811_C06 **(E)** and WT, *srf3-3*, *srf3-3* complementation line **(F)**. One-way ANOVA follows by a post-hoc Tukey HSD test, letters indicate statistical differences ($p<0.05$). **(G)** Graph of the late root growth response to low iron levels after 3 days provided by 100 µM, 50 µM or 10 µM of ferrozine and 300 µM of Na-Fe-EDTA on wild type (WT) and *srf3-2* and *srf3-4*. Independent two ways student test ($p<0.05$), n.s. non-significant. **(H)** Late root growth after transfer under standard condition in WT, *srf3-2*, *srf3-3* and *srf3-4*. One-way ANOVA follows by a post-hoc Tukey HSD test, letters indicate statistical differences ($p<0.05$). For boxplots, circles indicate a single biological replicate and the number below each box specifies the number of replicates, horizontal black bars indicate the median, the black cross represents the mean, box represents the interquartile range and the hinges the min and max whiskers.\

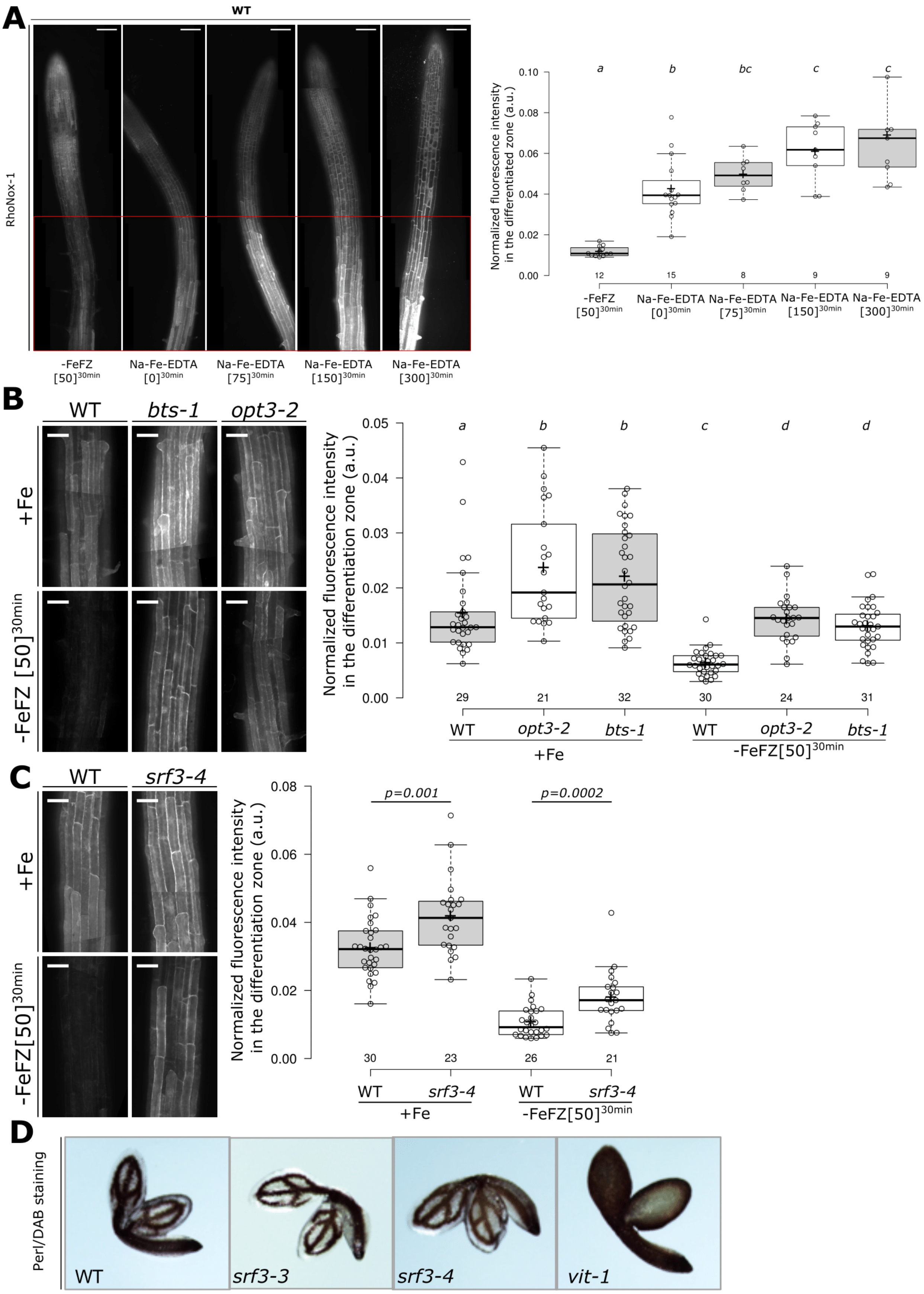
A
WT
RhoNox-1
-FeFZ [50]30min
Na-Fe-EDTA [0]30min
Na-Fe-EDTA [75]30min
Na-Fe-EDTA [150]30min
Na-Fe-EDTA [300]30min
Normalized fluorescence intensity in the differentiated zone (a.u.)
a
b
bc
c
c
12
15
8
9
9
B
WT
bts-1
opt3-2
+Fe
-FeFZ [50]30min
Normalized fluorescence intensity in the differentiation zone (a.u.)
a
b
b
c
d
d
29
21
32
30
24
31
WT
opt3-2
bts-1
WT
opt3-2
bts-1
+Fe
-FeFZ[50]30min
C
WT
srf3-4
+Fe
-FeFZ[50]30min
Normalized fluorescence intensity in the differentiation zone (a.u.)
p=0.001
p=0.0002
30
23
26
21
WT
srf3-4
WT
srf3-4
+Fe
-FeFZ[50]30min
D
Perl/DAB staining
WT
srf3-3
srf3-4
vit-1

**Supplementary Figure 2 (Related to Fig1). Characterization of RhoNox-1 staining and evaluation of iron content in embryo of WT, *srf3* mutants and *vit-1*. (A)** Left panel: confocal images of 5 days old seedling of WT epidermal root cells stained with RhoNox-1 pretreated with 50µM, 0 µM of FerroZine (FZ) and 75, 150 and 300 µM of Na-Fe-EDTA for 30 minutes. Red box indicates the region where the fluorescence intensity has been quantified. Scale bars, 100µm. Right panel: the related fluorescence intensity quantification. One-way ANOVA, follows by a post-hoc Tukey HSD test, letters indicate statistical differences ($p<0.05$). **(B)** Left panel: confocal images of differentiated epidermal root cells stained with RhoNox-1, WT, *bts-1* and *opt3-2* in sufficient (+Fe), upper panel, and in low iron levels (-Fe) with 50µM of ferrozine for 30 minutes. Scale bars, 50µm. Right panel: the related quantification of the normalized fluorescence intensity. One-way ANOVA, follows by a post-hoc Tukey HSD test, letters indicate statistical differences ($p<0.05$). **(C)** Left panel: confocal images of differentiated epidermal root cells stained with RhoNox-1, WT and *srf3-4* in sufficient (+Fe) and in low iron levels (-Fe) supplemented with 50µM of ferrozine for 30 minutes. Scale bars, 50µm. Right panel: related quantification of the normalized fluorescence intensity. One-way ANOVA followed by a post-hoc Tukey HSD test, letters indicate statistical differences ($p<0.05$). **(D)** Wild-type *srf3-3*, *srf3-4* and *vit-1* (positive control) dry seed embryos were dissected and stained with Perls/DAB method. For boxplots, circles indicate a single biological replicate and the number below each box specifies the number of replicates, horizontal black bars indicate the median, the black cross represents the mean, box represents the interquartile range and the hinges the min and max whiskers.

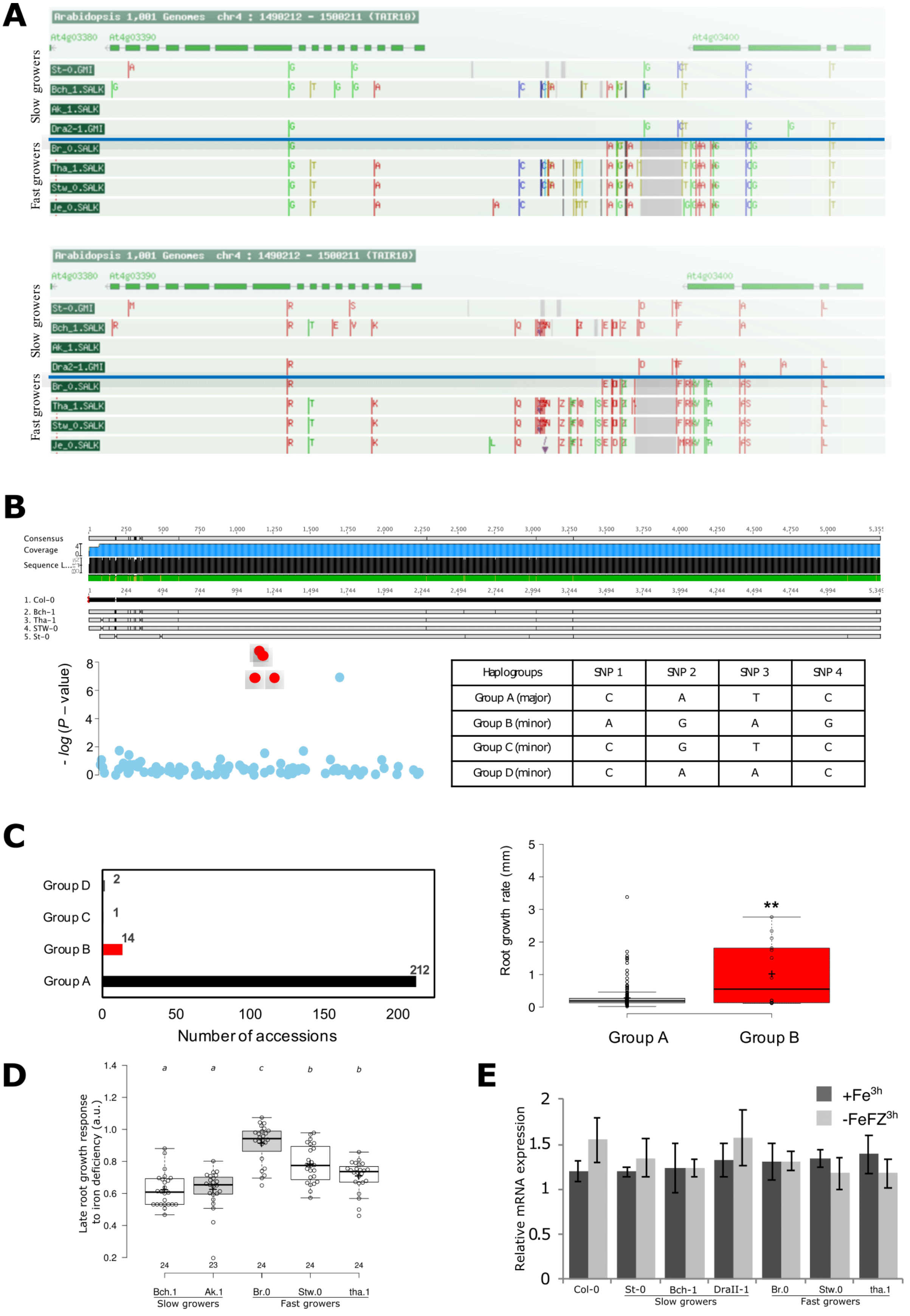
A
Arabidopsis 1,001 Genomes chr4 : 1490212 - 1500211 (TAIR10)
At4g03380
At4g03390
At4g03400
Slow growers
Fast growers
St-0.GMI
Bch_1.SALK
Ak_1.SALK
Dra2-1.GMI
Br_0.SALK
Tha_1.SALK
Stw_0.SALK
Je_0.SALK
B
Consensus
Coverage
1. Col-0
2. Bch-1
3. Tha-1
4. STW-0
5. St-0
- log (P – value)
Haplogroups | SNP 1 | SNP 2 | SNP 3 | SNP 4
Group A (major) | C | A | T | C
Group B (minor) | A | G | A | G
Group C (minor) | C | G | T | C
Group D (minor) | C | A | A | C
C
Group D
Group C
Group B
Group A
2
1
14
212
Number of accessions
Root growth rate (mm)
**
Group A
Group B
D
Late root growth response to iron deficiency (a.u.)
a
a
c
b
b
24
23
24
24
24
Bch.1
Ak.1
Br.0
Stw.0
tha.1
Slow growers
Fast growers
E
+Fe3h
-FeFZ3h
Relative mRNA expression
Col-0
St-0
Bch-1
DraII-1
Br.0
Stw.0
tha.1
Slow growers
Fast growers

**Supplementary Figure 3 (Related to Fig1). SNP polymorphism around the SRF3 locus and qPCR in extreme accessions**. **(A)** SNP polymorphism, top, and amino acid, bottom, changes surrounding the SRF3 locus in four representative accessions (http://signal.salk.edu/atg1001/3.0/gebrowser.php). **(B)** SNP polymorphisms in regulatory and coding regions of SRF3 gene in extreme accessions as confirmed by Sanger sequencing. SRF3 genomic DNA sequences from Bch-1, St-0, STW-0 and Tha-1 accessions were obtained by Sanger sequencing and aligned to Col-0 sequence and SNP changes (compared to Col-0 reference) from extreme accessions are shown in vertical lines. **(C)** Left panel: distribution of marker SNPs highlighted in red color in haplogroup A (major), B (minor), C (minor) and D (minor). Bar plot shows division of haplogroups A, B, C and D in 231 accessions. Right panel: box plots for root growth rate in Group A (major) and Group B (minor) accessions. Asterisks indicate significant difference with Tukey's HSD comparison (p-value < 0.05). **(D)** Late root growth response to low iron levels after transfer with extreme accession found in the GWAS. One-way ANOVA follows by a post-hoc Tukey HSD test, letters indicate statistical differences (p<0.05). **(E)** Relative expression level of SRF3 in Col-0, St-0, Bch-1, DraII-1, Br-0, Stw-0 and Tha-1 at T0, in mock treatment after 3 hours in low iron with 100 uM of ferrozine. For boxplots, circles indicate a single biological replicate and the number below each box specifies the number of replicates, horizontal black bars indicate the median, the black cross represents the mean, box represents the interquartile range and the hinges the min and max whiskers.

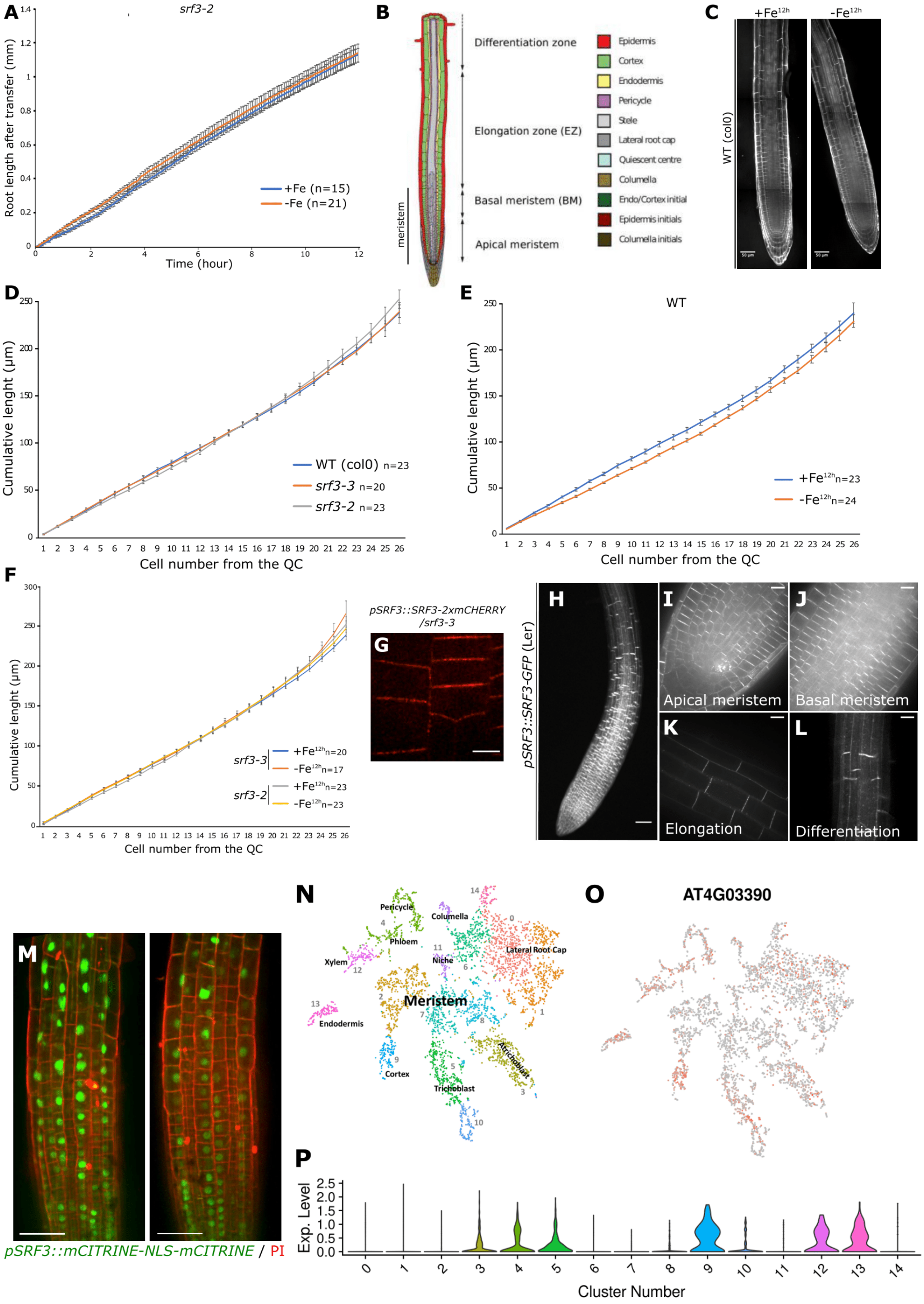
A
srf3-2
Root length after transfer (mm)
Time (hour)
+Fe (n=15)
-Fe (n=21)
B
Differentiation zone
Elongation zone (EZ)
Basal meristem (BM)
Apical meristem
meristem
Epidermis
Cortex
Endodermis
Pericycle
Stele
Lateral root cap
Quiescent centre
Columella
Endo/Cortex initial
Epidermis initials
Columella initials
C
+Fe12h
-Fe12h
WT (col0)
D
Cumulative lenght (µm)
WT (col0) n=23
srf3-3 n=20
srf3-2 n=23
Cell number from the QC
E
WT
+Fe12h n=23
-Fe12h n=24
F
srf3-3
srf3-2
+Fe12h n=20
-Fe12h n=17
+Fe12h n=23
-Fe12h n=23
pSRF3::SRF3-2xmCHERRY
/srf3-3
G
pSRF3::SRF3-GFP (Ler)
H
I
J
K
L
Apical meristem
Basal meristem
Elongation
Differentiation
M
pSRF3::mCITRINE-NLS-mCITRINE / PI
N
Pericycle
Columella
Phloem
Xylem
Lateral Root Cap
Niche
Meristem
Endodermis
Cortex
Trichoblast
Atrichoblast
O
AT4G03390
P
Exp. Level
Cluster Number

**Supplementary Figure 4 (Related to Fig2). SRF3 is transcription and translation co-exist in the transition-elongation zone and low iron triggers a root growth decrease in the elongation zone in an *SRF3*-dependent manner. (A)** Graph showing time lapse of the root length of WT and *srf3-2* under sufficient and low iron levels for 12 hours. Error bars indicated standard deviation of the mean (SEM). **(A)** Scheme of root tip showing the different zone of the root and the cell types. **(C)** Confocal images of 5 days old seedling stained with propidium iodide in WT under sufficient and low iron levels for 12 hours. Scale bars, 50µm. **(D)** Cumulative cell length from the quiescent center in WT under sufficient and low iron levels for 12 hours. Error bars indicated standard deviation of the mean (SEM). **(E)** Cumulative cell length from the quiescent center in WT under sufficient and low iron levels for 12 hours. Error bars indicated standard deviation of the mean (SEM). **(F)** Cumulative cell length from the quiescent center in *srf3* mutants under sufficient and low iron levels for 12 hours. Error bars indicated standard deviation of the mean (SEM). We observed that under low iron medium the roots present shorter cell length while the steepness of the curve representing the cumulative cell length was identical in both conditions in WT. However, no difference was noted in *srf3* mutants. This shows that the cell division process is not affected under low iron levels and that the decreased of the root growth is due to SRF3-dependent reduction of cell elongation. **(G)** Confocal image of root epidermal cells of plant expressing *pSRF3::SRF3-2xmCHERRY-4xmyc* in *srf3-3* (srf3-3 comp). Scale bar, 10µm. **(H, I, J, K,L)** Confocal images of 5 day-old seedlings expressing *pSRF3::SRF3-GFP* in root **(H)** (scale bar, 50µm), apical meristem **(I),** basal meristem **(J),** elongation **(K)** and differentiated tissue **(L)**. (Scale bars, 10µm). **(M)** Confocal images of 5 day-old seedlings expressing *pSRF3::mCITRINE-NLS-mCITRINE* and stain with propidium iodide (PI). Scale bars, 50µm. **(N)** Scheme of t-SNE plot represent gene expression across clusters in the different cell types. **(O)** t-SNE plots represent *SRF3* AT4G03390 gene expression across clusters in the different cell types. **(P)** Violin-plot depicting the distribution of expression levels for cells in the cluster. Y-axis (length) - gene expression level across each cluster. X-axis - proportion of cells showing a given expression value.

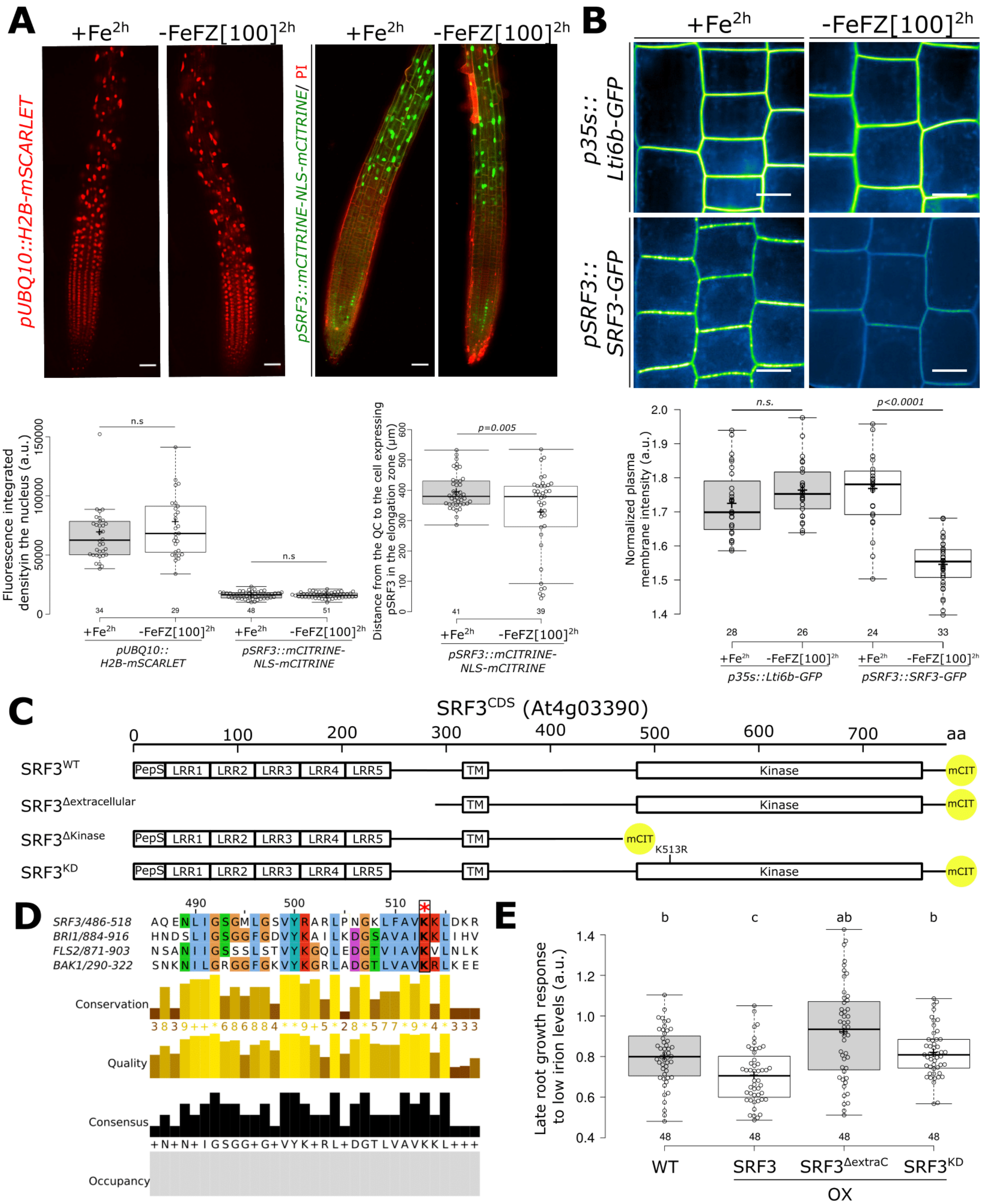


**Supplementary Figure 5 (Related to Fig2). The early lack of iron affects SRF3 protein levels at the plasma membrane which is dependent on the extracellular domain and kinase activity to regulate root growth. (A)** Confocal images of root tip of 5 days old seedling expressing *pUBQ10::H2B-mSCARLET* (Left) and *pSRF3::mCITRINE-NLS-mCITRINE* (right) with the plasma membrane stained by propidium iodide (PI) under mock (+Fe) and under low iron levels provided with 100 µM of ferrozine (-FeFZ[100]) for 2 hours. Scale bars, 50µm. Below, the related quantification of the nuclear fluorescence integrated density (lower left) and of the distance from the QC to the cell expressing pSRF3 in the elongation zone in µm (lower right). Independent two-ways student test ($p<0.05$), n.s. non-significant. (**B)** Left panel: Confocal images of root epidermis of 5 days old seedling expressing *p35s::Lti6b-GFP* and *pSRF3::SRF3-GFP* under mock (+Fe) and under low iron levels provided by 100 µM of ferrozine (FZ) for two hours. Scale bar, 10µm. Right panel: the related quantification of the normalized plasma membrane intensity. Independent two-ways student test ($p<0.05$), n.s. non-significant. **(C)** Scheme representing the SRF3 CDS full length as well as truncated version and mutated version. **(D)** Alignment done with Jalview of SRF3, BRI1, FLS2 and BAK1 ATP binding pocket. **(E)** Graph showing the quantification of late root growth response to low iron levels for WT and overexpressing lines of SRF3 CDS full length as well as truncated version and mutated version. One-way ANOVA followed by a post-hoc Tukey HSD test, letters indicate statistical differences ($p<0.05$). For boxplots, circles indicate a single biological replicate and the number below each box specifies the number of replicates, horizontal black bars indicate the median, the black cross represents the mean, box represents the interquartile range and the hinges the min and max whiskers.

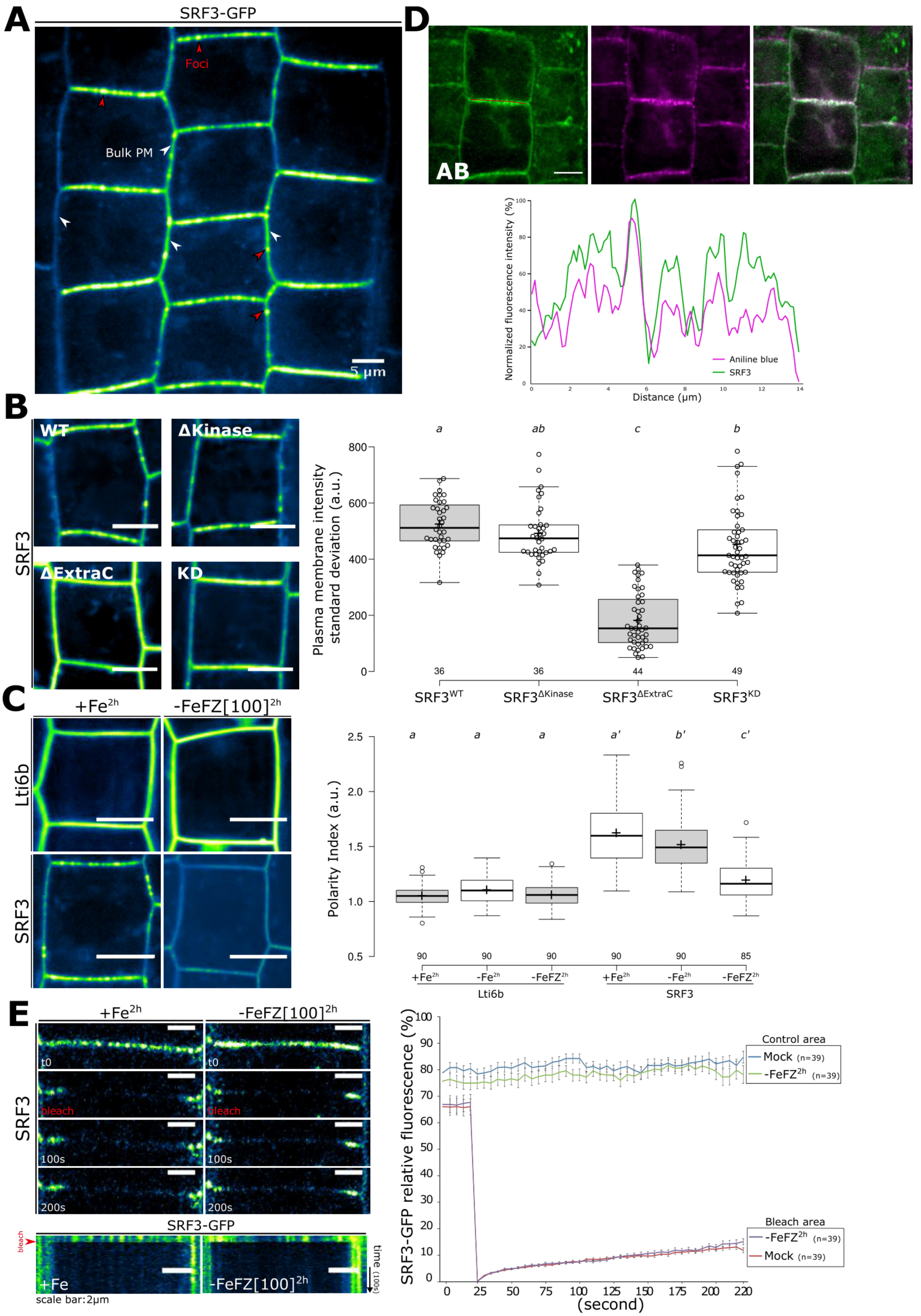
A
SRF3-GFP
Foci
Bulk PM
5 µm
D
AB
Normalized fluorescence intensity (%)
Distance (µm)
Aniline blue
SRF3
B
WT
ΔKinase
ΔExtraC
KD
SRF3
Plasma membrane intensity standard deviation (a.u.)
a
ab
c
b
36
36
44
49
SRF3WT
SRF3ΔKinase
SRF3ΔExtraC
SRF3KD
C
+Fe2h
-FeFZ[100]2h
Lti6b
SRF3
Polarity Index (a.u.)
a
a
a
a'
b'
c'
90
90
90
90
90
85
+Fe2h
-Fe2h
-FeFZ2h
Lti6b
SRF3
E
+Fe2h
-FeFZ[100]2h
t0
bleach
100s
200s
SRF3
SRF3-GFP
+Fe
-FeFZ[100]2h
time (100s)
scale bar:2µm
SRF3-GFP relative fluorescence (%)
Control area
Mock (n=39)
-FeFZ2h (n=39)
Bleach area
-FeFZ2h (n=39)
Mock (n=39)
(second)

**Supplementary Figure 6 (Related to Fig3). SRF3 is removed from the two PM-associated subpopulations under low iron. (A)** Confocal images of root epidermal cells in the elongation zone of 5 days old seedling expressing *pSRF3::SRF3-GFP*. Red arrows indicate foci and white arrows bulk plasma membrane (Bulk PM). Scale bar, 5µm. **(B)** Left panel: confocal images of root epidermis of 5 days old seedling expressing *pUBQ10::SRF3*$^{WT}$*-mCITRINE*, *pUBQ10::SRF3*$^{\Delta Kinase}$*-mCITRINE* ,*pUBQ10::SRF3*$^{\Delta ExtraC}$*-mCITRINE,* *pUBQ10::SRF3*$^{KD}$*-mCITRINE* (SRF3$^{WT}$, SRF3$^{\Delta Kinae}$, SRF3$^{\Delta ExtraC}$ and SRF3$^{KD}$). Scale bars, 10µm. Right panel: related quantification of the mean standard deviation of the intensity mean at the apical-basal side of the cell. One-way ANOVA followed by a post-hoc Tukey HSD test, letters indicate statistical differences ($p<0.05$). **(C)** Left panel: Confocal images of root epidermal cells in the elongation zone of 5 days old seedling expressing *p35s::Lti6b-GFP* (Lti6b) and *pSRF3::SRF3-GFP* (SRF3) under mock (+Fe) and under low iron media provided or not by 100 µM of ferrozine (-FeFZ[100]) for two hours. Scale bar, 5µm. Right panel related quantification of polarity index. One-way ANOVA followed by a post-hoc Tukey HSD test, letters indicate statistical differences ($p<0.05$). **(D)** Plant stained with the callose maker Aniline Blue (AB; left) and SRF3 (middle) and the relative merge (right), graph at the bottom indicates the signal intensity in both channel on the apical basal part of the cell. Red line on the left image indicates where the scan line has been traced. Scale bars, 10µm. **(E)** Left panel: confocal images of *pSRF3::SRF3-GFP* (SRF3) of 5 days old seedling during FRAP experiment in WT in the mock and under low iron media supplemented with 100µM of ferrozine and the related kymograph (time scale 15 seconds) on the bottom. Scale bar, 2µm. Right panel: traces of *pSRF3::SRF3-GFP* fluorescence intensity at the plasma membrane during FRAP analyses in the different conditions. For boxplots, circles indicate a single biological replicate and the number below each box specifies the number of replicates, horizontal black bars indicate the median, the black cross represents the mean, box represents the interquartile range and the hinges the min and max whiskers.

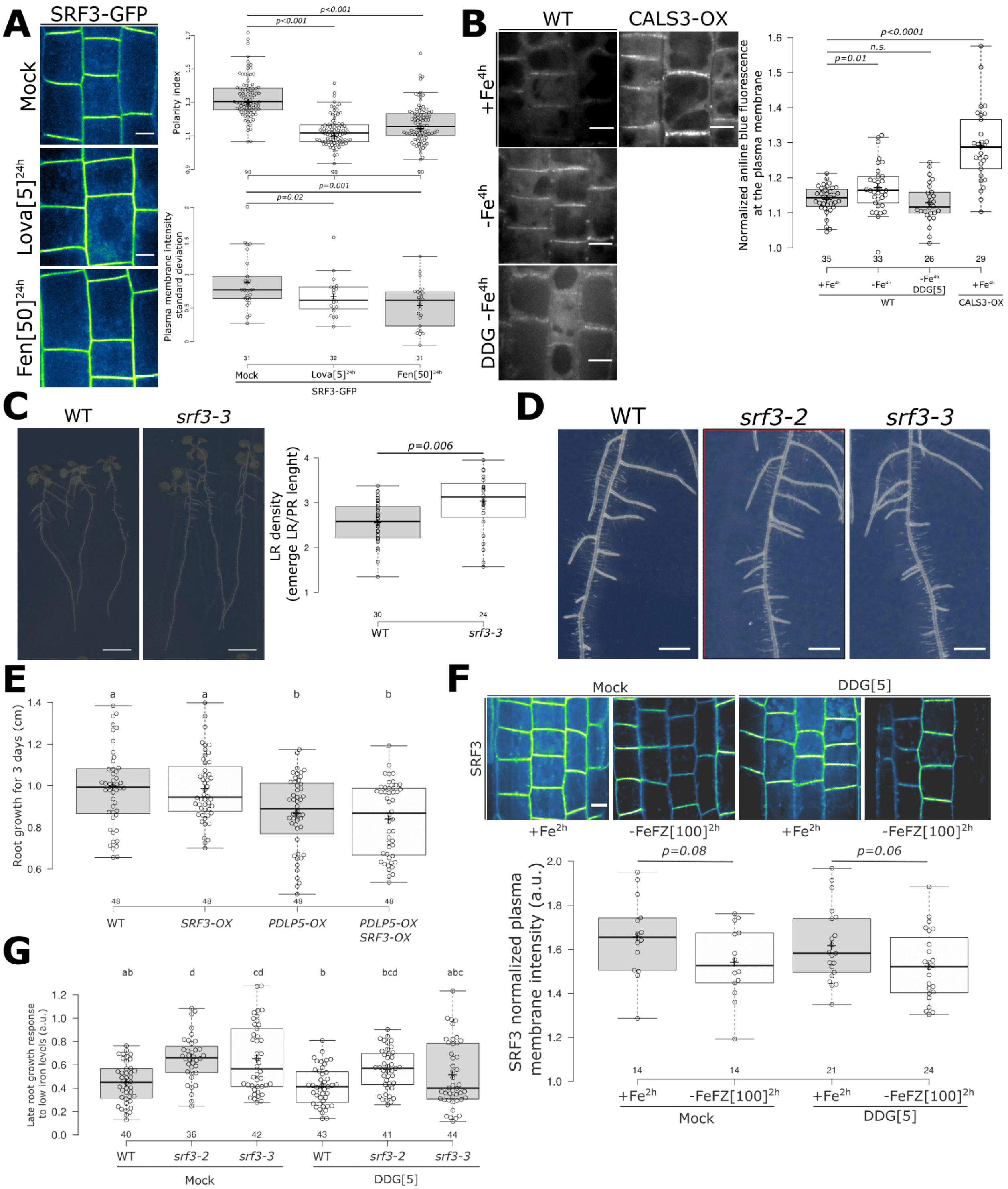


**Supplementary Figure 7 (Related to Fig4 & 5). SRF3 is an upstream negative regulator of callose synthases. (A)** Upper panel: confocal images of root epidermis of 5 days old seedling expressing *pSRF3::SRF3-GFP* under mock and treated for 24 hours with Lovastin (Lova) and Fenpropimorph (Fen). Scale bar, 10µm. Lower panel: related quantification of the polarity index and the standard deviation of the mean intensity at the apical basal part of the cell. **(B)** Left panel: confocal images of root epidermal cells in the elongation zone of 5 days old seedling stained with aniline blue (AB) in the indicated genotypes, WT and 35s::GFP-CALS3 under mock (+Fe) and under low iron media for four hours with or without 2-deoxy-d-glucose (DDG) Scale bar, 10µm. Right panel: the related quantification of normalized plasma membrane intensity. Independent two-ways student test ($p<0.05$), n.s. non-significant. **(C)** Upper panel: images of 12 days old seedling of WT and *srf3-3* under normal growth condition. Scale bar, 1cm. Lower panel: the quantification of the lateral root (LR) density. (**D**) Images of 12 days old seedling of WT and *srf3-2* and *srf3-3.* Scale bar, 2mm. **(E)** Graph representing the quantification of the mean root growth rate for 3 days WT, *UBQ10::SRF3-mCITRINE* (*SRF3-OX), 35s::PDLP5-GFP (PDLP5-OX)* and *PDLP5-GFPxSRF3-OX.* One-way ANOVA followed by a post-hoc Tukey HSD test, letters indicate statistical differences ($p<0.05$). **(F)** Upper panel: confocal images of root epidermis in the elongation zone of 5 days old seedling expressing *pUBQ10::SRF3-mCITRINE* in iron sufficient and deficient media supplemented with 100uM of ferrozine for 2 hours in presence or absence of DDG. Scale bar, 10µm. Lower panel: related quantification of the normalized plasma membrane intensity. Independent two-ways student test ($p<0.05$). **(G)** Graph representing the root growth response to low iron levels for 3 days with or without 2-deoxy-d-glucose (DDG) in WT, *srf3-2* and *srf3-3.* One-way ANOVA followed by a post-hoc Tukey HSD test, letters indicate statistical differences ($p<0.05$). For boxplots, circles indicate a single biological replicate and the number below each box specifies the number of replicates, horizontal black bars indicate the median, the black cross represents the mean, box represents the interquartile range and the hinges the min and max whiskers.

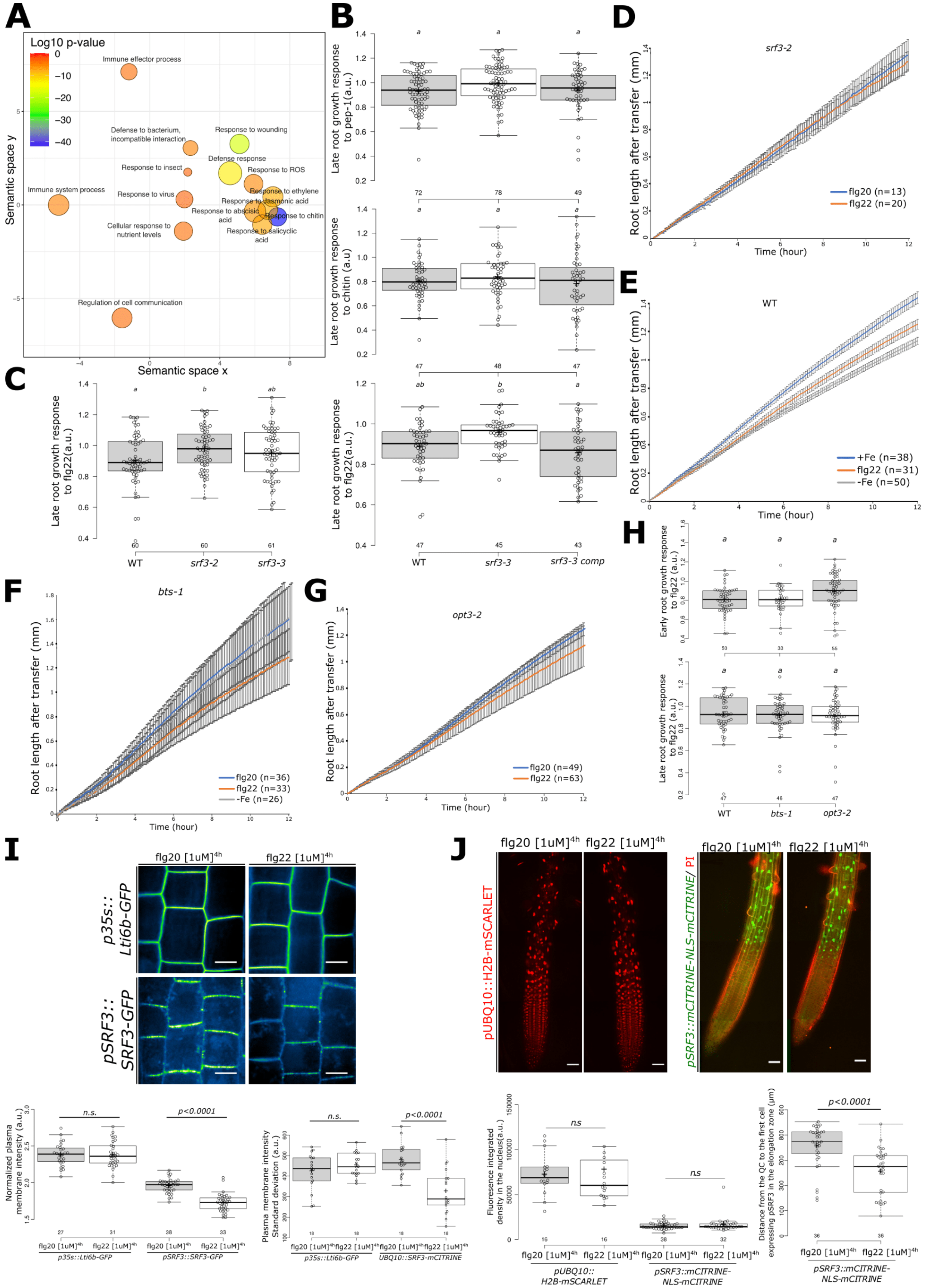
A
Log10 p-value
Immune effector process
Defense to bacterium, incompatible interaction
Response to wounding
Defense response
Response to insect
Response to ROS
Immune system process
Response to virus
Response to ethylene
Response to jasmonic acid
Response to abscisic acid
Response to chitin
Cellular response to nutrient levels
Response to salicylic acid
Regulation of cell communication
Semantic space x
Semantic space y
B
Late root growth response to pep-1(a.u.)
Late root growth response to chitin (a.u)
Late root growth response to flg22(a.u.)
WT
srf3-3
srf3-3 comp
C
Late root growth response to flg22(a.u.)
WT
srf3-2
srf3-3
D
srf3-2
Root length after transfer (mm)
Time (hour)
flg20 (n=13)
flg22 (n=20)
E
WT
+Fe (n=38)
flg22 (n=31)
-Fe (n=50)
F
bts-1
flg20 (n=36)
flg22 (n=33)
-Fe (n=26)
G
opt3-2
flg20 (n=49)
flg22 (n=63)
H
Early root growth response to flg22 (a.u.)
Late root growth response to flg22 (a.u.)
WT
bts-1
opt3-2
I
flg20 [1uM]4h
flg22 [1uM]4h
p35s::Lti6b-GFP
pSRF3::SRF3-GFP
J
pUBQ10::H2B-mSCARLET
pSRF3::mCITRINE-NLS-mCITRINE/ PI
Normalized plasma membrane intensity (a.u.)
n.s.
p<0.0001
Plasma membrane intensity Standard deviation (a.u.)
UBQ10::SRF3-mCITRINE
Fluoresence integrated density in the nucleus(a.u.)
ns
pUBQ10:: H2B-mSCARLET
pSRF3::mCITRINE-NLS-mCITRINE
Distance from the QC to the first cell expressing pSRF3 in the elongation zone (µm)

**Supplementary Figure 8 (Related to Fig6). Specific regulation of flg22-induced bacterial root innate immunity by SRF3. (A)** Gene ontology of differential expressed genes in srf3 mutants compare to the WT under normal growth condition. **(B)** Box plot showing the late root growth response to pep-1, top panel, chitin, middle panel and flg22, bottom panel in WT, *srf3-3*, *srf3-3* complementation. One-way ANOVA followed by a post-hoc Tukey HSD test, letters indicate statistical differences (p<0.05). (**C)** Box plot showing the late root growth response to flg22 in WT, *srf3*-2 and *srf3-3*. One-way ANOVA followed by a post-hoc Tukey HSD test, letters indicate statistical differences (p<0.05). (**D)** Graph showing time lapse of the root length of WT and *srf3-2* under flg20 and flg22 for 12 hours. Error bars indicated standard deviation of the mean (SEM). **(E)** Graph showing time lapse of the root length of WT under iron sufficient and low iron media and media supplemented with flg22 at 1µM for 12 hours**.** Error bars indicated standard deviation of the mean (SEM). **(F)** Graph showing time lapse of the root length of *bts-1* under flg20 and flg22 at 1µM for 12 hours. **(G)** Graph showing time lapse of the root length of *opt3-2* under iron deficiency, flg20 and flg22 at 1µM for 12 hours. Error bars indicated standard deviation of the mean (SEM). **(H)** Box plot of the early, top panel, and late, bottom panel, root growth response to flg22 of WT, *fls2-c*, *bts-1* and *opt3-2* for 12 hours and 3 days. One-way ANOVA followed by a post-hoc Tukey HSD test, letters indicate statistical differences (p<0.05). **(I)** Top panel: Confocal images of root epidermis of 5 days old seedling expressing *p35s::Lti6b-GFP* and *pSRF3::SRF3-GFP* (*Ler* background) under flg20 and flg22 at 1µM for four hours. Scale bars, 10µm. Bottom left panel: related quantification of the normalized plasma membrane intensity. Bottom right panel: quantification of the standard deviation of the signal intensity at the apical basal side of the PM in *p35s::Lti6b-GFP* and *pUBQ10::SRF3-mCITRINE* under flg20 and flg22 at 1µM for four hours. Independent two-ways student test (p<0.05), n.s. non-significant. **(J)** Upper panel: confocal images of root tip of 5 days old seedling expressing *pUBQ10::H2B-mSCARLET* and *pSRF3::mCITRINE-NLS*-mCITRINE with the plasma membrane stained by propidium iodide (PI) under flg22 at 1µM for four hours, scale bars, 50µm, the related quantification of the nuclear fluorescence integrated density (bottom left) and of the distance from the QC to the cell expressing pSRF3 in the elongation zone in µm (bottom right). Independent two-ways student test (p<0.05), n.s. non-significant. For boxplots, circles indicate a single biological replicate and the number below each box specifies the number of replicates, horizontal black bars indicate the median, the black cross represents the mean, box represents the interquartile range and the hinges the min and max whiskers.

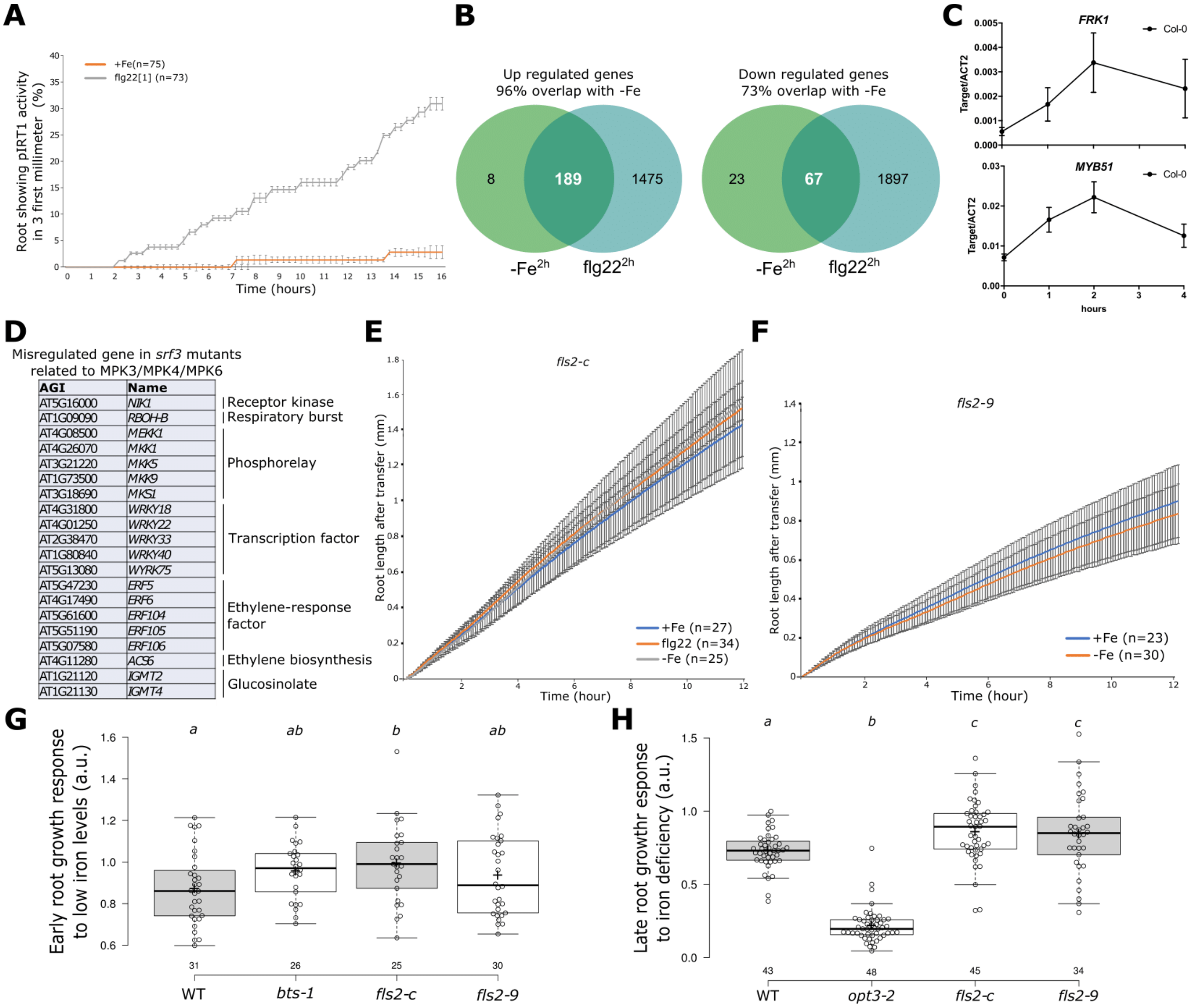


**Supplementary Figure 9 (Related to Fig6). SRF3 is involved in PTI signaling pathways regulation. (A)** Quantification of time lapse analysis of pIRT1::NLS-2xYPet under mock and flg22 at 1µM treatment for 16 hours. **(B)** Venn diagram of differentially expressed genes, up regulated and down regulated, under iron deficiency and flg22 for 2 hours. **(C)** Graph representing the expression level of immune-related genes, *FRK1* top and *MYB51* bottom under low iron levels according to time. Error bars indicate SEM. **(D)** List of PIT-dependent genes mis regulated in *srf3* mutants belonging to the early iron/defense core regulatory network. **(E, F)** Graph showing time lapse of the root length of WT under sufficient (+Fe) and low (-Fe) iron levels media and flg22 at 1uM for 12 hours in *fls2-c* **(E)** and *fls2-9* mutants **(F), (G)** the related quantification including bts-1 mutant as a positive control. One-way ANOVA followed by a post-hoc Tukey HSD test, letters indicate statistical differences ($p<0.05$). **(H)** Quantification of the late root growth response to iron deficiency media in *fls2-c* and *fls2-9* mutants and opt3-2 used as a positive control. One-way ANOVA followed by a post-hoc Tukey HSD test, letters indicate statistical differences ($p<0.05$). For boxplots, circles indicate a single biological replicate and the number below each box specifies the number of replicates, horizontal black bars indicate the median, the black cross represents the mean, box represents the interquartile range and the hinges the min and max whiskers.
